# The solid tumor microenvironment changes the hierarchy of CD155 and CD112 receptors, shaping checkpoint blockade outcome

**DOI:** 10.64898/2026.07.29.741502

**Authors:** Valentina Carannante, Karl Olofsson, Hanqing Zhang, Niklas Sandström, Jacopo Fontana, Damien Toullec, Grace Turyasingura, Birte Hell, Arnika Kathleen Wagner, Patrick A. Sandoz, Hanna Van Ooijen, Andreas Lundqvist, Martin Wiklund, Björn Önfelt

**Affiliations:** Dept. of Microbiology, Tumor and Cell biology, Science for Life Laboratory, Karolinska Institutet, 171 65 Solna, Stockholm, Sweden; Dept. of Applied Physics, Science for Life Laboratory, KTH Royal Institute of Technology, 171 65 Solna, Stockholm, Sweden; Dept. of Medicine, Karolinska Institutet, 171 76 Stockholm, Sweden; Dept. of Oncology-Pathology, Karolinska Institutet, 171 65 Solna, Stockholm, Sweden

## Abstract

Reproducing a physiologically relevant tumor microenvironment *in vitro* is essential for developing effective immunotherapeutic treatments. By integrating the use of combinatorial receptor blockade and organoid models we provide a deep functional understanding of CD155 and CD112 receptors in solid tumors and their impact on cellular immunotherapy and infiltration. CD226 showed plasticity in response to the environment, being able to switch between CD155 and CD112 depending on the ligand availability. In addition, CD226 drove NK cell infiltration into tumor tissues via CD155 and CD112 ligation, with CD226-CD112 interaction specifically promoting migration from the periphery to the core. Downregulation of CD155 and TIGIT induced by the tumor microenvironment and previous drug exposure reduced the long-term efficacy of TIGIT blockade. Taken together, our findings point towards using CD112R blockade in primary tumors to simultaneously enhance NK cell killing activity and promote infiltration into the tumor core via CD226 and CD112 interaction.

**ONE SENTENCE SUMMARY:** Tumors shape the hierarchy of CD155-CD112 receptors, reducing TIGIT blockade efficacy, while CD226 drives NK infiltration and shows binding plasticity

## INTRODUCTION

Immune checkpoint inhibitors (ICIs) is an established clinical approach to treat cancer by restoring immune cell antitumoral activity and proliferative capacity, inducing durable responses in patients (*1*). However, a large subset of cancer patients does not respond to ICIs, because of innate or acquired resistance to therapy (*2*). Today, the main challenge is to reduce the number of cancer patients not responding to ICIs, finding new molecular targets and combining already existing therapeutic approaches (*3*). In this context, the interest in Nectin-like molecules CD155 and CD112 has dramatically increased in recent years. Nectin-like molecules carry out multiple functions in human biology, regulating cell proliferation (*4*), cell motility and adhesion (*5*), angiogenesis (*6*) and immune cell recognition (*7–12*). As dysregulation of these functions leads to tumor transformation (*13*), it is not surprising that CD155 and CD112 are overexpressed in several cancers, correlating positively with tumor burden and metastatic progression (*14–16*), and negatively with disease-free survival rate (*16*). Ongoing clinical trials are focused on directly targeting CD155 with oncolytic viral therapy, or indirectly with the use of anti-TIGIT and anti-CD112R antibodies.

TIGIT and CD112R are inhibitory receptors binding to CD155 and CD112 (*8*, *10*, *17*). They are preferentially expressed on natural killer (NK) cells and T cells and can be upregulated in cancer patients (*18–21*), correlating with poor prognosis (*22*). There are currently more than 40 ongoing clinical trials targeting the TIGIT pathway. In 2020, promising results were obtained in the phase II CITYSCAPE trial with the combination of anti-TIGIT antibody (tiragolumab) with anti-PDL1 (atezolizumab) for the treatment of stage IV non-small cell lung cancer patients (NSCLC) with high PD-L1 expression (*23*). However, the follow-up phase III trial SKYSCRAPER-01 did not confirm the benefits (*24*). These data highlight the importance of better understanding the role and regulation of CD155 and CD112 in cancer to help predict which patients that may benefit from TIGIT and CD112R blockade.

NK cells are NKp46^+^ innate lymphocytes responsible for initiating and sustaining anti-tumoral immune responses (*25*). Therapeutic strategies that enhance NK cell activity are considered promising alternatives for patients resistant to T cell-based treatments (*26*). The activity of NK cells is finely regulated by a balance of activating and inhibitory signals (*27*), and the main aim of ICI therapy, such as anti-TIGIT and anti-CD112R treatment, is to dampen inhibition and shift the balance toward activation. However, it is not possible to understand the impact of TIGIT blockade without considering the role of the other CD155 receptors, CD226 (*7*, *28*), CD96 (*11*, *29*) and KIR2DL5 (*12*, *30*) co-expressed on NK cells.

CD226 is an activating CD155-receptor that promotes NK and T cell-mediated cytotoxicity and cytokine release (*28*, *31*). CD226 can also interact with CD112 with lower binding affinity compared to CD155 (*32*). Despite the activating role of CD226, CD155-deficient mice show reduced tumor growth and metastasis (*33*). It is therefore unclear to what extent CD226 activation relies on CD155 interaction, and what the impact of such interaction is on NK cell-mediated tumor surveillance and migration.

CD96 is a single-pass transmembrane glycoprotein mostly expressed on NK and T cells (*11*), which levels peak between 6 to 9 days after activation on T cells (*29*). CD96 binding to CD155 has been shown to reduce IFN-γ secretion in murine NK cells (*34*), and its blockade enhanced NK cell-mediated surveillance of tumor metastasis in mouse models (*35*).

KIR2DL5 (CD158f) is an inhibitory receptor belonging to human KIR family (*30*, *36*), and it was recently shown to bind CD155 (*12*). It is expressed in a small fraction of CD56^Dim^ NK and T cells (*30*, *36*, *37*), and only in some individuals (*30*).

Thus, CD155 and CD112-mediated regulation of NK cells is intricate, where TIGIT, CD96, KIR2DL5, CD226 and CD112R compete for their ligands, and the outcome in part depends on which interaction prevails. Since such outcome might have a direct impact on the efficacy of checkpoint blockade therapy, better understanding of the contribution from each CD155-CD112 receptors in modulating NK cell activity in the tumor microenvironment is needed.

In this paper, we study the hierarchy of CD155 and CD112 receptors in modulating NK cell degranulation, cytotoxic activity and infiltration in the tumor microenvironment. To increase the translational impact of our findings, we performed flow cytometry and live cell imaging assays in both traditional 2D culture and newly developed 3D culture systems, the latter being more representative of the solid tumor microenvironment. Our study provides a better comprehension of the role of CD155 and CD112 in the treatment of tumors, and their implication in regulating NK cell tumor surveillance and the efficacy of immunotherapy.

## RESULTS

### The expression of CD155 inhibitory receptors TIGIT and KIR2DL5 is balanced in KIR2DL5^+^ donors

We examined the expression of CD226, CD112R, CD96, TIGIT, and KIR2DL5 on overnight activated NK cells (Fig. 1A, Fig. S1A, 1B). CD226 and CD112R were expressed by the majority of NK cells (Fig. 1A). CD96 was expressed in both CD56^Dim^ and CD56^Bright^ NK cells, although higher expression was observed in the CD56Bright subset (Fig. 1A, Fig. S1A). The expression of TIGIT was bimodal (Fig. 1A), with TIGIT^+^ NK cells mainly found in the CD56^Dim^ subset (Fig. S1B). KIR2DL5 is expressed only in a fraction of individuals (*30*) and only on a small fraction of the NK cells (Fig. 1B). Like TIGIT, the majority of the KIR2DL5^+^ cells were found in the CD56^Dim^ subset (Fig. 1B). As TIGIT and KIR2DL5 are expressed by the same NK cell subset and have overlapping function by giving inhibitory input upon CD155 interaction (*9*, *12*, *36*), we investigated if the expression patterns of these receptors were correlated (Fig. 1C-E). Indeed, higher levels of TIGIT were found in KIR2DL5^-^ NK cells compared to KIR2DL5^+^ NK cells, both in terms of frequency and mean fluorescence intensity (MFI) (Fig. 1D, 1E). These data show that the expression of CD155 inhibitory receptors on NK cells is diversified, ensuring recognition by both CD56^Bright^ and CD56^Dim^ NK cells via CD96 and TIGIT, respectively. Likewise, NK inhibition might be counterbalanced by reduced TIGIT levels in KIR2DL5^+^ cells, as a mechanism to maintain a consistent threshold of NK cell activation across various subsets in KIR2DL5^+^ donors. Such diversification was not observed for CD226 and CD112R.

**Fig. 1.**
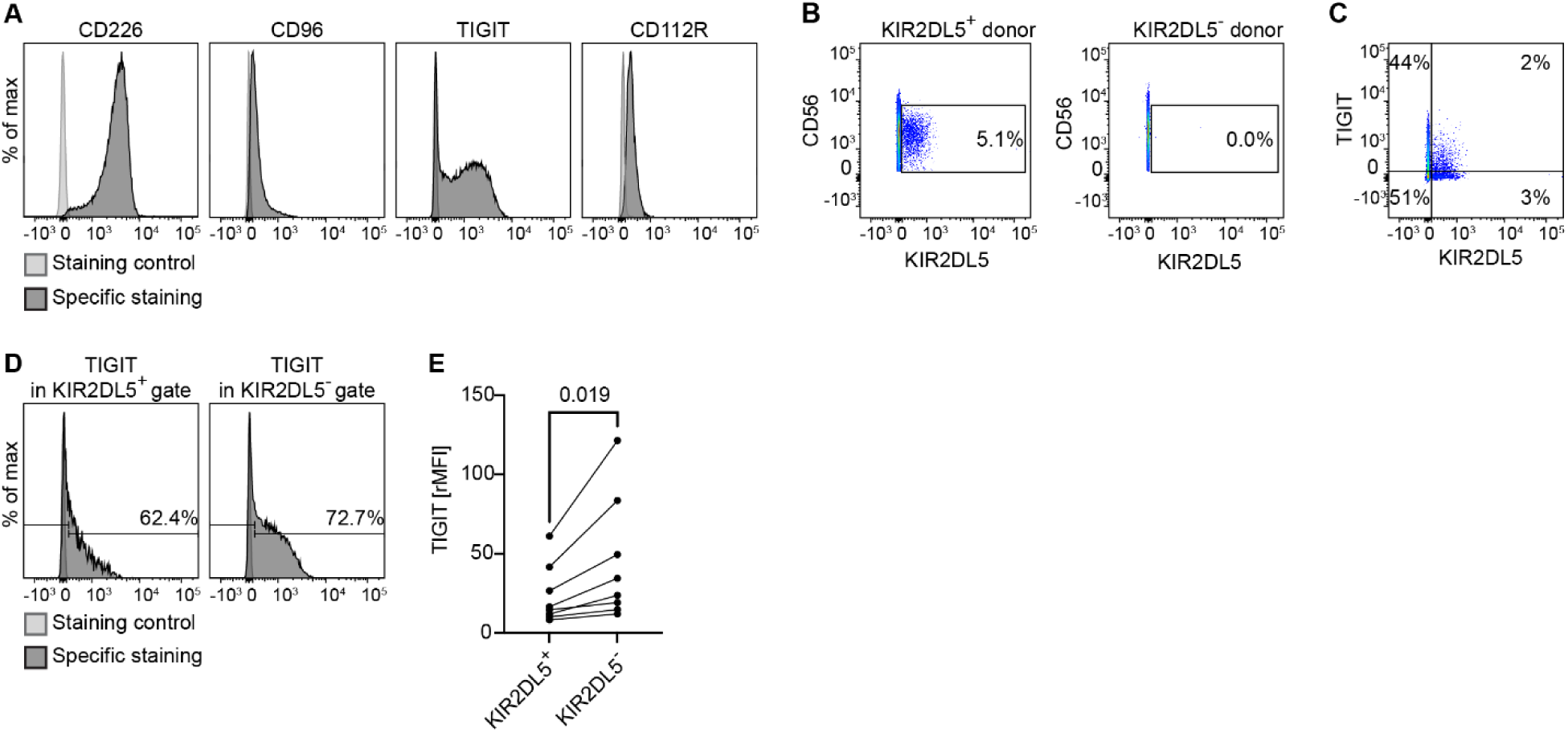
Expression of NK cells receptors of CD155 and CD112 at early stages of activation. **A.** A representative example of CD226, CD112R, CD96, TIGIT and KIR2DL5 expression on NK cells activated with IL-15 overnight. **B.** Flow cytometry dot-plots showing representative examples of KIR2DL5^+^ and KIR2DL5^-^ donors. **C.** Representative dot-plots showing the expression of TIGIT in relation to KIR2DL5 on overnight IL-15-activated NK cells isolated from a KIR2DL5^+^ donor. **D.** Representative dot-plots showing the expression of TIGIT in KIR2DL5^+^ (left panel) and KIR2DL5^-^ (right panel) NK cells isolated from a KIR2LD5^+^ donor. **E.** Pairwise comparison indicating the relative MFI of TIGIT in the KIR2DL5^+^ and KIR2DL5^-^ NK cells gates. Only KIR2LD5^+^ donors were considered in the analysis. Statistical method: paired t-test (n=8).

### NK cell response to CD155 in renal carcinoma cells is dominated by TIGIT- and KIR2DL5-mediated inhibition, and CD226 shows plasticity in response to ligand modulation

To characterize the impact of each CD155 and CD112 receptor and their interplay on NK recognition, we used the renal carcinoma cell line A498 which expresses high levels of the two ligands and generated single and double knock-out versions for CD155 and CD112 (Fig. 2A, Fig.S1C). Then, we exposed NK cells to the different A498 cell variants and assessed degranulation by measuring CD107a surface expression. Opposite responses were observed in TIGIT^+^ and TIGIT^-^ NK cells: TIGIT^+^ NK cells showed enhanced degranulation against single knock-out A498 ^CD155-/-CD112wt^ and double knock-out A498 ^CD155-/-CD112-/-^ cells (Fig.2B), while suppressed activity was observed in TIGIT^-^ NK cells in the same conditions (Fig.2C). No differences were observed when both TIGIT^+^ and TIGIT^-^ NK cells were exposed to single knock-out A498 ^CD155wt^ ^CD112-/-^ cells (Fig.2B, 2C). Therefore, CD155 expression leads to different behaviors in TIGIT^+^ and TIGIT^-^ NK populations. As TIGIT^-^ NK cells express CD226, these data suggest a predominant effect of TIGIT-mediated signaling over CD226 in TIGIT^+^ NK cells.

**Fig. 2.**
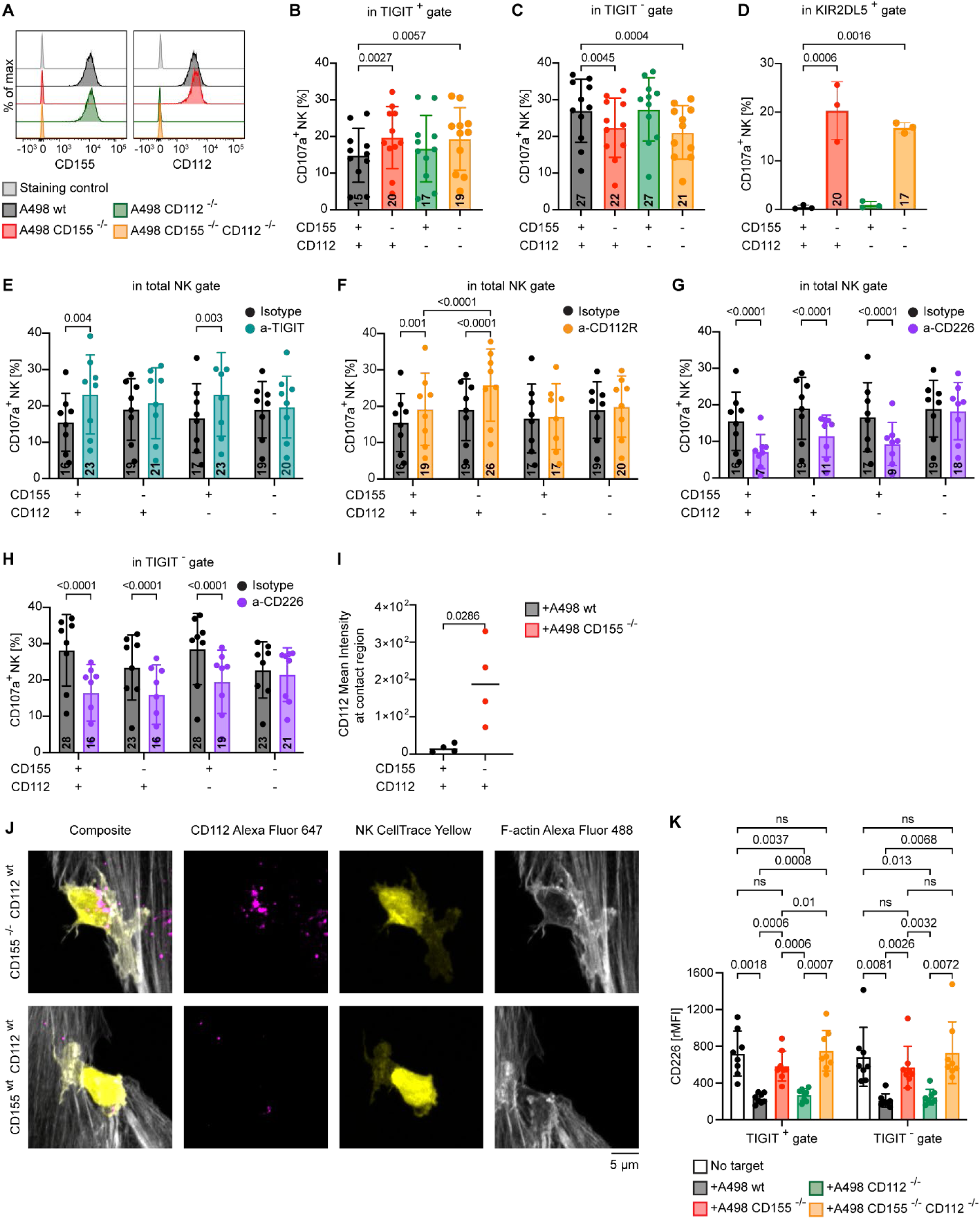
TIGIT and KIR2DL5-mediated inhibition constitute the primary response to CD155, while CD226 contribution to NK cell degranulation depends on both CD155 and CD112. **A.** CD155 and CD112 surface expression on A498 ^CD155wt^ ^CD112wt^ (black histogram), A498^CD155-/-CD112wt^ (red histogram), A498^CD155wt^ ^CD112-/-^ (green histogram) and A498^CD155-/-CD112-/-^ (orange histogram). **B, C.** Percentage of degranulating NK cells in response to A498 ^CD155wt^ ^CD112wt^ (black bar), A498^CD155-/-CD112wt^ (red bar), A498^CD155wt^ ^CD112-/-^ (green bar) and A498^CD155-/-CD112-/-^ (orange bar) in TIGIT^+^ (**B**) TIGIT^-^ (**C**) gates (n=8). **D.** Percentage of degranulating NK cells in KIR2DL5^+^ gate in KIR2DL5^+^ donors in response to A498wt (black bar), A498^CD155-/-CD112wt^ (red bar), A498^CD155wt^ ^CD112-/-^ (green bar) and A498^CD155-/-CD112-/-^ (orange bar). **E, F, G.** Effect of TIGIT blockade (teal bars), CD112R blockade (orange bars), and CD226 blockade (purple bars) on NK cell degranulation after exposure to A498 ^CD15wt CD112wt^ (+,+), A498^CD155-/- CD112wt^ (-,+), A498CD155wt CD112-/- (+,-) and A498CD155 -/- CD112-/- (-,-) as indicated on the x-axis. **H.** Effect of CD226 blockade (purple) on degranulation in TIGIT^-^ NK cells exposed to A498^CD155wt CD112wt^ (+,+), A498^CD155-/- CD112wt^ (-,+), A498^CD155wt CD112-/-^ (+,-) and A498^CD155 -/- CD112-/-^ (-,-) as indicated on the x-axis. **I.** Mean intensity fluorescence of CD112 in the region of contact between NK cells and A498^CD155wt^ ^CD112wt^ (black bar) or A498^CD155-/-CD112wt^ (red bar) (n=4). **J.** Representative images showing CD112 (in magenta) recruited in the region of contact between NK cells (in yellow) and A498^CD155-/-CD112wt^ (upper panel) versus A498^CD155wt^ ^CD112wt^ (lower panel). F-actin is in green. **K.** CD226 surface expression on NK cells after exposure to A498wt (black bar), A498^CD155-/-CD112wt^ (red bar), A498^CD155wt^ ^CD112-/-^ (green bar) and A498^CD155-/-CD112-/-^ (orange bar) compared to no target control (white bar) in TIGIT^+^ gate (left panel) and TIGIT^-^ gate (right panel) (n=8). The numbers within the bars in graphs B to H represent the percentage of NK cels positive for CD107a. Statistical tests: One-way ANOVA followed by Dunnet’s multiple comparison test was performed in B-D. Two-way ANOVA followed by Tukeýs post hoc test was performed in E-H.

A similar pattern was observed for KIR2DL5^+^ and KIR2DL5^-^ NK cells in KIR2DL5^+^ donors (Fig.2D, Fig.1SD-F), further reinforcing the notion that inhibitory receptors for CD155 overcome CD226-mediated signaling. Interesting, the loss of CD112 did not significantly affect NK cell degranulation (Fig.2B-D, Fig.1SD-G), suggesting that TIGIT, KIR2DL5 and CD226 preferentially bind to CD155 rather than CD112.

CD112 has been recently shown to interact with the inhibitory receptor CD112R on NK cells, potentially affecting NK cell responses in our experimental conditions. To characterize CD112R impact on the TIGIT, KIR2DL5 and CD226 axis, we performed the same set of experiments in the presence of blocking antibodies for TIGIT, CD112R and CD226 (Fig.2E-G). TIGIT blockade improved NK cell degranulation only when NK cells were exposed to A498 ^CD155wt^ ^CD112wt^ and A498 ^CD155wt^ ^CD112-/-^ cells (Fig.2E), confirming that TIGIT mainly ligates to CD155 but not to CD112. CD112R blockade was effective against A498 ^CD155wt^ ^CD112wt^ and A498 ^CD155-/-CD112wt^, but not against A498 ^CD155wt^ ^CD112-/-^, confirming the specificity of CD112R for CD112 (Fig.2F). In the presence of CD112R blocking antibody, NK cells showed higher activation against A498 ^CD155-/-CD112wt^ compared to A498 ^CD155wt^ ^CD112wt^ (Fig.2F), indicating that simultaneous loss of CD155 and CD112 signaling through TIGIT and CD112R might have an additive effect on NK cell anti-tumor activity. This hypothesis was further reinforced by the evidence that improved degranulation was exclusively observed in TIGIT^+^ NK cells, and not in TIGIT^-^ NK cells (Fig.1SH, 1SI). A similar trend was observed in KIR2DL5^+^ TIGIT^-^ and KIR2DL5^-^ TIGIT^-^ NK cells (Fig.1SJ, 1SK). Therefore, TIGIT and KIR2DL5 bind to CD155 and CD112R to CD112, and the combination of CD155 loss and CD112R receptor blockade leads to enhanced degranulation in TIGIT^+^ and KIR2DL5^+^ NK cells.

The pattern of preferential binding for a single ligand was interrupted when we performed CD226 blocking experiments. As expected, CD226 blockade impaired NK cell degranulation against A498 ^CD155wt^ ^CD112wt^, as well as against A498 ^CD155-/-CD112wt^ and A498 ^CD155wt^ ^CD112-/-^, but not in the absence of both ligands (A498 ^CD155-/-CD112-/-^) (Fig.2G), confirming that CD226 binds to both CD155 and CD112. To exclude any effect caused by TIGIT, we gated in TIGIT^-^ NK cells (Fig.2H), corroborating the previous findings.

Next, we asked if CD226 binds to both CD155 and CD112 at steady state or if it preferentially binds to one of the two ligands. Given the higher affinity of CD226 for CD155 compared to CD112, we hypothesized that CD226 preferentially binds to CD155 and switches to CD112 in its absence. The absence of differences in NK degranulation between A498 ^CD155wt^ ^CD112wt^ and A498 ^CD155-/-CD112wt^ while blocking CD226 also suggested this hypothesis (Fig.2H). We looked in the region of contact between A498 cells and NK cells to verify the presence of CD112 by high-resolution microscopy (Fig.2I, 2J). We observed CD112 accumulation in the contact between NK cells and A498 ^CD155-/-CD112wt^ cells (Fig.2I, Fig.2J upper panel), but not in the A498 ^CD155wt^ ^CD112wt^ control (Fig.2I, Fig.2J lower panel). These results confirmed that CD226 preferentially binds to CD155, and it switches to CD112 in the absence of CD155.

Last, we wanted to investigate if CD226 competes with TIGIT and KIR2DL5 for CD155 binding. As CD155 interaction induces CD226 down-modulation, we measured CD226 expression in TIGIT^+^ NK cells and TIGIT^-^ NK after exposure to A498 variants (Fig.2K). As expected, CD226 down-modulation occurred after exposure to CD155 in A498 ^CD155wt^ ^CD112wt^ and A498 ^CD155wt^ ^CD112-/-^ conditions, but no significant differences were observed between TIGIT^+^ and TIGIT^-^ NK (Fig.2K, Fig.1SL), indicating that CD226 interacts with CD155 at similar extent in these two populations. On the other hand, differences were observed between KIR2DL5^+^ and KIR2DL5^-^ NK cells, suggesting that competition for CD155 might occur between CD226 and KIR2DL5 in KIR2DL5^+^ donors (Fig.1SM). Interestingly, CD112 interaction did not induce CD226 down-regulation (Fig.2K). and no differences in CD226 expression were observed when blocking CD112R in our experimental settings (Fig.S1N).

In summary, TIGIT, KIR2DL5 and CD226 preferentially bind to CD155 rather than CD112, with TIGIT and KIR2DL5 overriding CD226-mediated signaling. For KIR2DL5 this might be the result of direct ligand binding competition, while no evidence for this was found for TIGIT. CD112R binds specifically to CD112, and simultaneous CD155 loss and CD112R receptor blockade leads to enhanced degranulation in TIGIT^+^ and KIR2DL5^+^ NK cells. While inhibitory receptors tend to bind to either CD155 or CD112, CD226 shows more plasticity in response to the environment, being able to switch between CD155 and CD112 depending on the ligand-availability on tumor cells.

### Variations of CD155 expression on renal carcinoma cells are sensed by TIGIT and KIR2DL5, while CD226 plasticity reduces the dependency on CD155 expression

Next, we investigated how NK cell recognition was affected by variations in CD155-expression on target cells. By performing siRNA-mediated CD155 knockdown on A498 ^CD155wt^ ^CD112wt^ for different time periods (24-96 hours) we obtained a wide range of CD155 expression levels (Fig. 3A, B). The expression of CD155 decreased with incubation time, reaching approximately 10% of that of A498 ^CD155wt^ ^CD112wt^ at the final timepoint (96 hours) (Fig. 3A, B). CD155 expression remained unvaried on A498 ^CD155wt^ ^CD112wt^ treated with scramble siRNA (Fig. 3B, Fig. S2A). In addition, siRNA-induced CD155 modulation was specific, as the expression of other NK cell ligands on A498 ^CD155wt^ ^CD112wt^ remained unchanged during treatment (Fig. S2B).

**Fig. 3.**
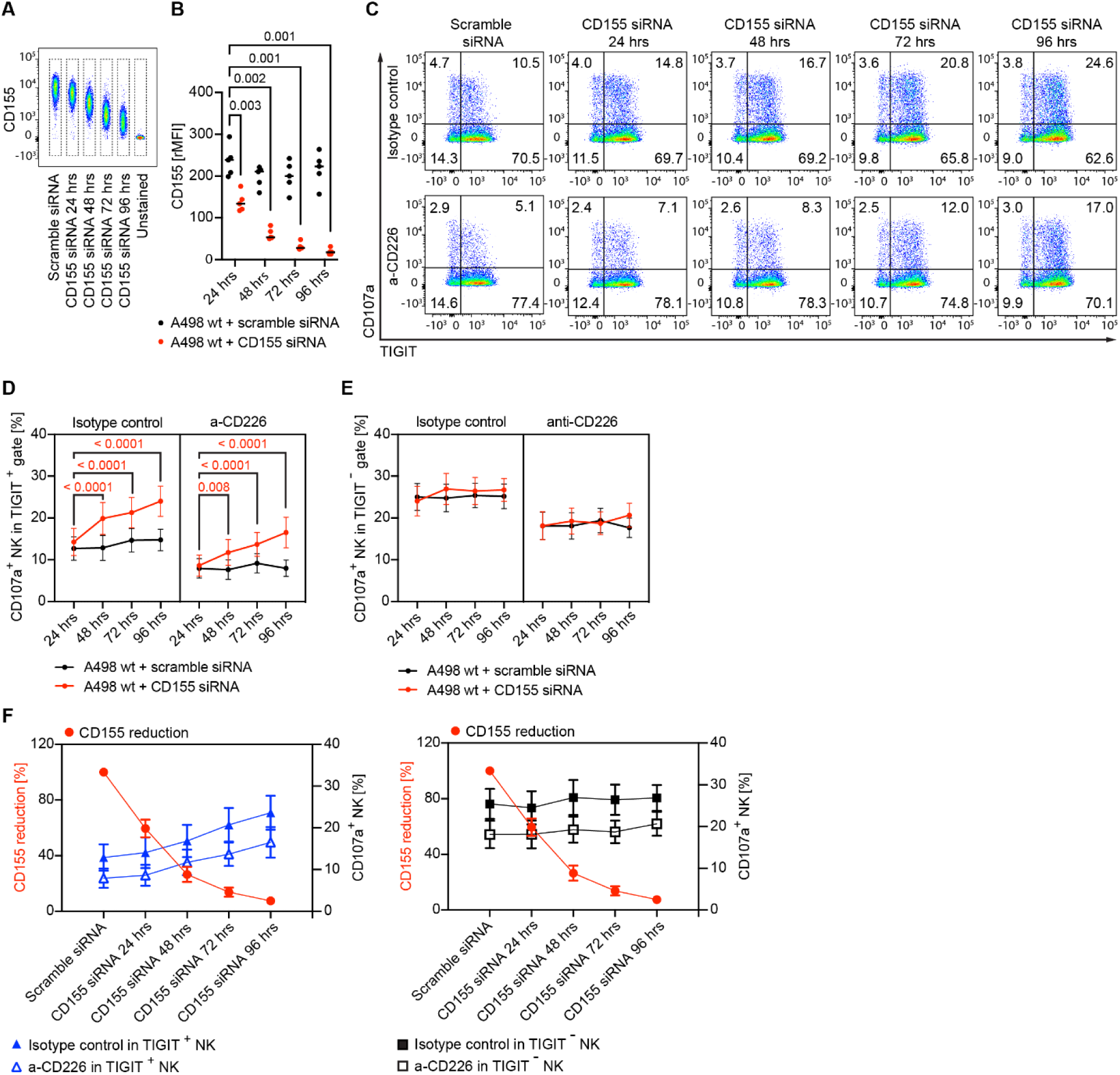
Varied CD155 expression on renal carcinoma tumor cells is sensed by TIGIT but not CD226 A, **B.** CD155 expression on A498wt cells incubated with either scramble siRNA (black in **B**) or CD155 siRNA (red in **B**) for 24, 48, 72 or 96 hours. In **B**: statistical method: two-way ANOVA followed by Tukey’s Post Hoc test (n=5). The black lines represent median values. **C.** NK degranulation (CD107a^+^) in relation to TIGIT expression. Overnight IL-15 activated NK cells were incubated with A498wt pre-treated with either scramble siRNA or CD155 si-RNA, in presence of an isotype control antibody (top row) or anti-CD226 blocking antibody (bottom row). **D. E.** Percentage of degranulating TIGIT^+^ (**D**) or TIGIT^-^ (**E**) NK cells in response to A498wt pre-treated with either scramble siRNA (black) or CD155 si-RNA (red), in the presence of an isotype control antibody (left) or anti-CD226 (right). Statistical method: two-way ANOVA followed by Dunnett’s Post Hoc test (n=8). The dots represent mean values and the lines represent standard error of the mean (SEM). Mean +/- SD values are reported in Supporting Table 1. **F.** Summarizing graphs of NK cell recognition of CD155 expression on renal carcinoma tumor cells. CD155 reduction (red dots), calculated by dividing the CD155 relative MFI of each condition by the CD155 relative MFI of the control (scramble siRNA at 24 hours). Left graph, in blue: CD107a^+^ TIGIT^+^ NK cells in presence of anti-CD226 (empty triangles) or isotype control antibody (filled triangles). Right graph, in black: CD107a^+^ TIGIT^-^ NK cells in presence of anti-CD226 (empty squares) or isotype control antibody (filled squares).

Degranulation assays with overnight IL-15-activated NK cells targeting A498 ^CD155wt^ ^CD112wt^ cells treated with either CD155-siRNA or scramble-siRNA for different times showed that the percentage of degranulating NK cells increased in response to decreasing CD155 expression, while no variations were observed in the control condition (Fig. S2C). The response was specifically induced by CD155 expression variations, as no differences in CD107a degranulation were observed against A498 ^CD155-/-CD112wt^ cells that had been treated with CD155-siRNA for different times (Fig. S2D, E). By gating on TIGIT^+/-^ NK cells it became evident that the increased degranulation against A498 ^CD155wt^ ^CD112wt^ with decreasing levels of CD155 was driven by TIGIT^+^ NK cells (Fig. 3C-E). In addition, the variations in response to CD155-siRNA treatment for different time periods disappeared in the presence of TIGIT blocking antibody (Fig. S2F).

In KIR2DL5^+^ donors, the fraction of degranulating KIR2DL5^+^ TIGIT^-^ NK cells increased with reduced CD155 expression (Fig. S2G, H), although the contribution of KIR2DL5^+^ NK cells to the total NK degranulation was minimal (Fig. S2G). These experiments further confirmed that alterations in CD155-expression modulated NK cell degranulation primarly via TIGIT and KIR2DL5 rather than CD226.

CD226 blockade reduced the overall NK cell degranulation but an activation in response to decreasing CD155-levels was still observed (Fig. 3C, D, Fig. S2C). We then looked specifically at TIGIT^-^ NK cells to understand how CD226 senses variations in CD155-expression. CD155 downmodulation did not alter NK cell degranulation in the TIGIT^-^ gates (Fig. 3E), suggesting that CD226 switches to CD112 not only in the absence of CD155, but also when CD155 levels decrease. The biological meaning of CD226 down-regulation induced by CD155 and not CD112 combined with its plasticity in response to ligand viability, and its impact on NK cell serial killing remains to be defined.

Figure 3F summarizes the responses of TIGIT^+^ and TIGIT^-^ NK cells to A498 renal carcinoma cells with different levels of CD155-expression. Overall, TIGIT^-^ NK cells respond stronger than TIGIT^+^ NK cells, but only TIGIT^+^ NK cells respond to the variation in CD155 expression. Blocking CD226 has a significant effect on both NK cell subsets in terms of degranulation, but the amplitude of this effect is independent of CD155 levels on the target cell, further reinforcing the evidence of CD226 switching to CD112 in the absence or low levels of CD155.

### Spheroid models of renal carcinoma cells to study NK cell activity in the tumor microenvironment

The hierarchy of CD155 and CD112 receptor activation described in our model suggested that TIGIT blockade should improve NK-mediated tumor surveillance if a consistent pool of TIGIT^+^ NK cells and high levels of CD155 are present intratumorally. To test this hypothesis, we studied the hierarchy of CD155 and CD112 receptors in renal carcinoma spheroids. We obtained renal carcinoma spheroids using the ultrasonic standing wave (USW)-assisted formation developed by our group (Fig.4A) (*38*). As control for some experiments, tumor spheroids with similar shape and size were formed in standard agarose microwells (*39*) (Fig.S3A, B).

**Fig. 4.**
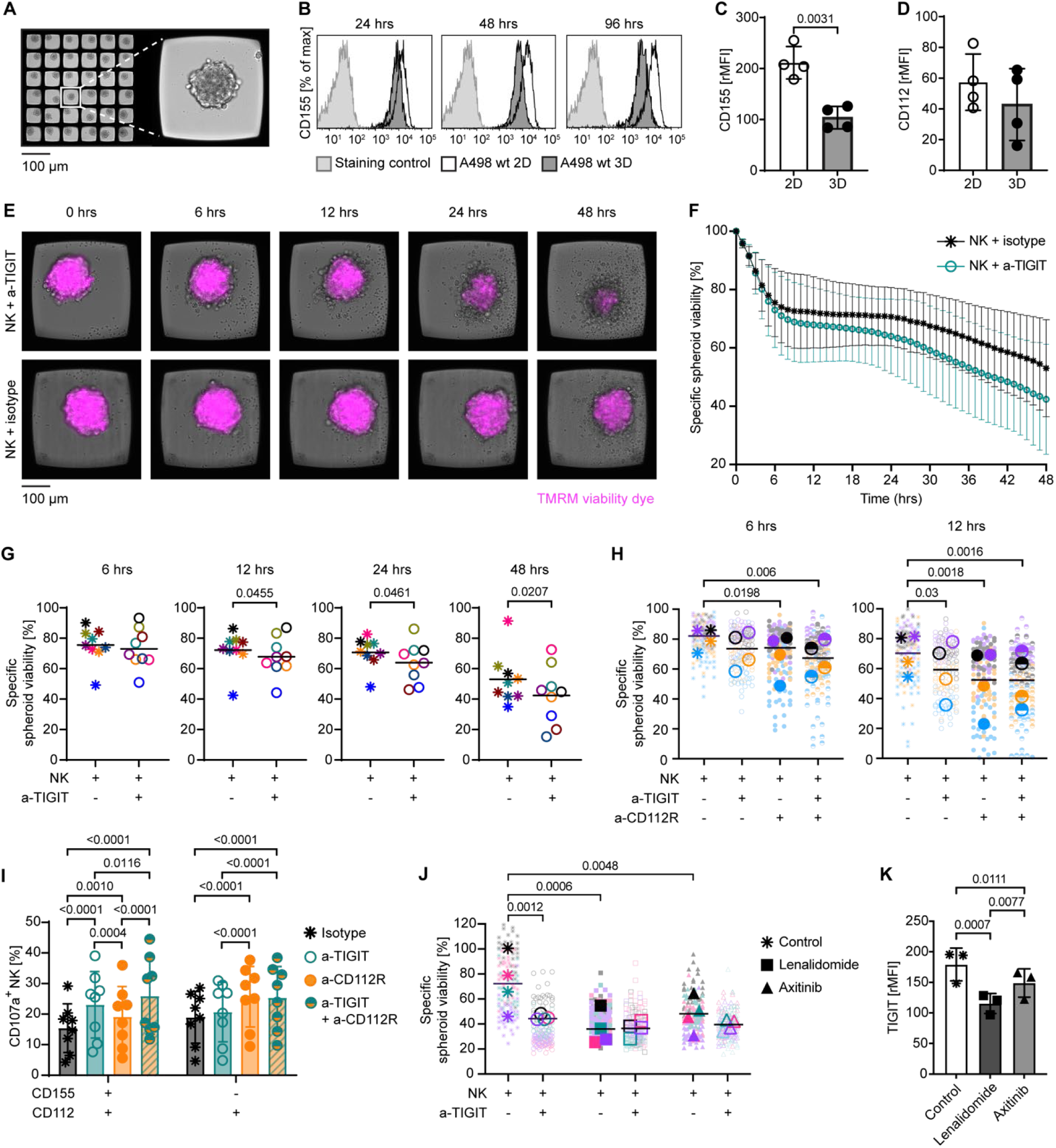
Implications of TIGIT expression on NK cell responses against A498 renal carcinoma spheroids. **A.** USW-induced A498 renal carcinoma spheroids obtained in the multichambered microwell chip (t=48 hours). Left panel: representative image of a single microwell chip chamber, containing 36 USW-induced spheroids sharing the same culture medium. Right panel: representative image of a single USW-induced spheroid in a microwell. **B.** CD155 expression on A498 ^CD155wt^ ^CD112wt^ cells maintained either as monolayers (white) or spheroids (dark grey) for 24, 48 or 96 hours. **C, D.** Statistical analysis of CD155 (**C**) and CD112 (**D**) expression in A498 ^CD155wt^ ^CD112wt^ cells maintained either as monolayers (white) or spheroids (dark grey) for 48 hours. Statistical method: unpaired t-test (n=4). **E.** Time-lapses of NK-mediated spheroid killing in presence of anti-TIGIT antibody (upper panel) or isotype control antibody (bottom panel). NK cells were pre-activated overnight with IL-15. In magenta: TMRM viability dye. Effector to target E:T seeding ratio = 1:1. **F.** Time-course of specific viability of A498 ^CD155wt^ ^CD112wt^ spheroids treated with NK cells in the presence of isotype control (black stars) or anti-TIGIT (teal empty dots). Mean values and SD of the nine donors for each condition are shown. **G.** Statistical analysis of specific viability of A498 ^CD155wt^ ^CD112wt^ spheroids with NK cells in the presence of isotype control (stars) or anti-TIGIT (empty dots) at different timepoints. The big symbols represent the mean values for 36 spheroids; each color represents different donor. Statistical test: paired t-test (n=9). **H.** Statistical analysis of specific viability of A498 ^CD155wt^ ^CD112wt^ spheroids at 6 hours (left panel) and 12 hours (right panel) with NK cells in the presence of isotype control (stars), anti-TIGIT (empty dots), anti-CD112R (filled dots) and anti-TIGIT + anti-CD112R (half filled dots). The big symbols represent the mean values for 36 spheroids; each color represents different donor. Statistical test: one-way ANOVA followed by Tukey’s Post Hoc test (n=4). **I.** Percentage of degranulating NK cells in response to A498 ^CD155wt^ ^CD112wt^ (left) or A498^CD155-/-CD112wt^ (right) in the presence of anti-TIGIT (empty dots, teal), anti-CD112R (filled dots, orange) and anti-TIGIT + anti-CD112R (half-filled dots, teal and orange). Statistical test: two-way ANOVA followed by Tukey’s Post Hoc test (n=8). **J.** Statistical analysis of specific viability of A498 ^CD155wt^ ^CD112wt^ spheroids with NK cells in the presence of isotype control (stars), lenalidomide (squares) or axitinib (triangles) in the presence (empty shapes) or in the absence (filled shapes) of anti-TIGIT. The big symbols represent the mean values for 36 spheroids; each color represents different donor. Statistical test: two-way ANOVA followed by Tukey’s Post Hoc test (n=4). **K.** Effect of lenalidomide (squares) or axitinib (triangles) on TIGIT expression of NK cells cultured in the presence of IL-15 (t= 48 hours). Statistical method: one-way ANOVA followed by Tukey’s Post Hoc test (n=3).

CD155 downregulation was observed in tumor spheroids over time (Fig. 4B, Fig.S3C), corresponding to a 20% and 50% reduction at 24 and 48 hours respectively (Fig.S3D). CD155 expression levels were stabilized after 48 hours and maintained up to 96 hours (Fig.4B, Fig.S3C). Stable tumor spheroid conformation was usually reached after around 48 hours (*40*), suggesting that maturation of cell-to-cell contacts could be involved in regulating the level of CD155 expression in solid tissues. As cell viability was similar between A498 ^CD155wt^ ^CD112wt^ cells grown as monolayers and as tumor spheroids (Fig.S3E), and the CD155 expression levels were independent of the 3D culture method used (Fig.S3F), we concluded that lower levels of CD155 expression were a property of the 3D solid tumor architecture. On the other hand, no statistical difference was observed in CD112 expression in tumor spheroids compared to 2D cultures (Fig.4D, Fig.S3G). In summary, lower levels of CD155 and similar levels of CD112 were available for NK cell recognition in tumor spheroids compared to standard tumor monolayers. Tumor spheroids were enzymatically dissociated and incubated with IL15-activated NK cells. In agreement with our previous results, NK cells responded better to A498 ^CD155wt^ ^CD112wt^ cells isolated from tumor spheroids (CD155^low^ cells) compared to 2D cultures (CD155^high^ cells) (Fig.S3H), with differences mainly observed in TIGIT^+^ NK cells (Fig.S3I, S3J). TIGIT blockade increased the percentage of CD107a^+^ NK cells, indicating that the levels of CD155, even if low, were enough to inhibit TIGIT^+^ NK cell degranulation (Fig.S3K).

To evaluate the effect of the solid tumor microenvironment on TIGIT therapy, we tested the efficacy of TIGIT blockade on NK cells directly exposed to tumor spheroids. Figure 4E shows an example of NK cell-mediated A498 ^CD155wt^ ^CD112wt^ renal carcinoma spheroid killing in the multi-chambered microwell chip over time with or without TIGIT blockade. TIGIT blockade significantly enhanced the ability of NK cells to kill A498 ^CD155wt^ ^CD112wt^ spheroids as indicated by the decreasing levels of TMRM signal and changes of spheroid texture over time (Fig.4E). The impact of TIGIT blockade on NK killing was delayed compared to cell monolayers, with the first effect being detected at 6 hours but statistical significance reached at 12 hours (Fig. 4F, 4G) and maintained up to the end of our assay (Fig. 4E-G). TIGIT blocking antibody had no effect on the viability of A498 ^CD155wt^ ^CD112wt^ spheroids in the absence of NK cells (Fig.S3L, S3M).

TIGIT blockade did not induce a complete spheroid eradication, with residual live tumor cells mainly localized in the spheroid core. The analysis of TIGIT expression on NK cells retrieved from tumor spheroids confirmed the infiltration of TIGIT^+^ NK cells in the tissue (Fig.S3N). However, both percentage of TIGIT^+^ NK cells and TIGIT expression were lower in tumor spheroids compared to NK maintained in suspension (Fig.S3N, S3O). Therefore, low CD155 availability combined with decreased expression of TIGIT might reduce the impact of TIGIT inhibition of NK cell killing in solid tumors, potentially affecting TIGIT blockade long-term efficacy and causing treatment failure.

As CD112 expression was stable in renal carcinoma spheroids, we investigated whether the combination of CD112R and TIGIT blockade could accelerate NK activity in solid tumors. Indeed, CD112R blockade improved NK killing against A498 ^CD155wt^ ^CD112wt^ spheroids at 6 hours, when TIGIT blockade did not show efficacy (Fig.4H, left panel). Both TIGIT and CD112R blockade improved NK killing against A498 ^CD155wt^ ^CD112wt^ spheroids at 12 hours, with a tendency of CD112R blockade of performing better in some donors, but no further effects were observed using combinatorial treatment (Fig.4H, right panel). When exposing NK cells to A498 ^CD155wt^ ^CD112wt^ monolayers (Fig.4I), both TIGIT and CD112R blockade improved NK degranulation, with TIGIT blockade having a predominant effect, and combinatorial TIGIT and CD112R blockade did improve the overall degranulation (Fig.4I). On the other hand, CD112R blockade had a major impact in NK cell degranulation against A498 ^CD155-/-CD112wt^, and combinatorial treatment did not lead to further improvements (Fig.4I). Therefore, our data indicates that the solid tumor microenvironment alters the dominance balance between TIGIT and CD112R due to less CD155 availability and lower TIGIT expression.

Last, we tested TIGIT blockade in combination with lenalidomide and axitinib. Lenalidomide is an immunomodulatory drug recently proposed for treatment of refractory metastatic renal carcinoma (*41*). Axitinib is a second-generation vascular endothelial growth factor-receptor (VEGF-R) tyrosine kinase inhibitor (TKI) used for treatment of advanced renal carcinoma (*42*). In the absence of NK cells, axitinib decreased the A498 ^CD155wt^ ^CD112wt^ spheroid viability to 93%, while lenalidomide had no effect (Fig.S3P).

Combining lenalidomide or axitinib with NK cells decreased the spheroid viability to 36% and 48% respectively from the 72% observed for NK cells alone (Fig. 4J). No improvements were observed when combining anti-TIGIT with lenalidomide or axitinib (Fig. 4J). Treating NK cells with IL-15 with axitinib or lenalidomide reduced the expression of TIGIT (Fig. 4K), explaining the absence of improvements using combinatorial treatment. These data further reinforce the importance of receptor modulation induced by the tumor microenvironment and drug exposure on TIGIT efficacy against solid tumors. This data further reinforces the importance of ligand and receptor modulation induced by the tumor microenvironment and drug exposure on TIGIT blockade efficacy against solid tumors. These results also point toward the use of CD112R blockade as a safer option due to less modulation of CD112 by the solid tumor architecture.

### CD226 promotes NK cell infiltration into the tumor core via CD112 ligation

In our previous experiments, viable tumor cells could still be observed in the spheroid core at the end of NK cell 3D killing assays, raising the question of whether NK cells could penetrate the spheroid core already at early timepoints and what would be the involvement of CD226 plasticity on NK overall cell migration. To answer these questions, we developed a novel imaging-based infiltration assay and an automated 3D analysis pipeline quantifying NK presence in different spheroid areas.

IL-15 activated NK cells were incubated with A498 ^CD155wt^ ^CD112wt^ for 5 hours (Fig.S3Q), followed by deep-tissue imaging and 3D segmentation to identify the tumor mass (in cyan) and tumor-infiltrating NK cells (in red) (Fig.5A, B). Changes in spheroid texture and size could be induced by NK killing, affecting the interpretation of NK cell infiltration data. The time point “5 hours” was identified as optimal, as no changes in spheroid radii were observed compared to the initial measurement (Fig.S3Q). To characterize NK infiltration, we analyzed the distribution of NK^+^ volume from the spheroid surface (Fig.5C) and quantified the area under the curve at intervals of 10 μm (Fig.5D), approximately corresponding to a single tumor cell layer. The analysis in A498 ^CD155wt^ ^CD112wt^ spheroids showed 44% of NK volume distributed within 10 μm from the surface (Fig.5C, D). The remaining NK cells that were able to infiltrate the first tumor barrier mainly localized in the region between 10 to 20 μm (33% of NK volume), with 15% of NK volume detected in the region between 20 and 30 μm, and only 8% of the NK volume detected beyond 30 μm from the spheroid surface. Thus, a minority of NK cells were able to infiltrate beyond two layers of tumor cells.

**Fig. 5.**
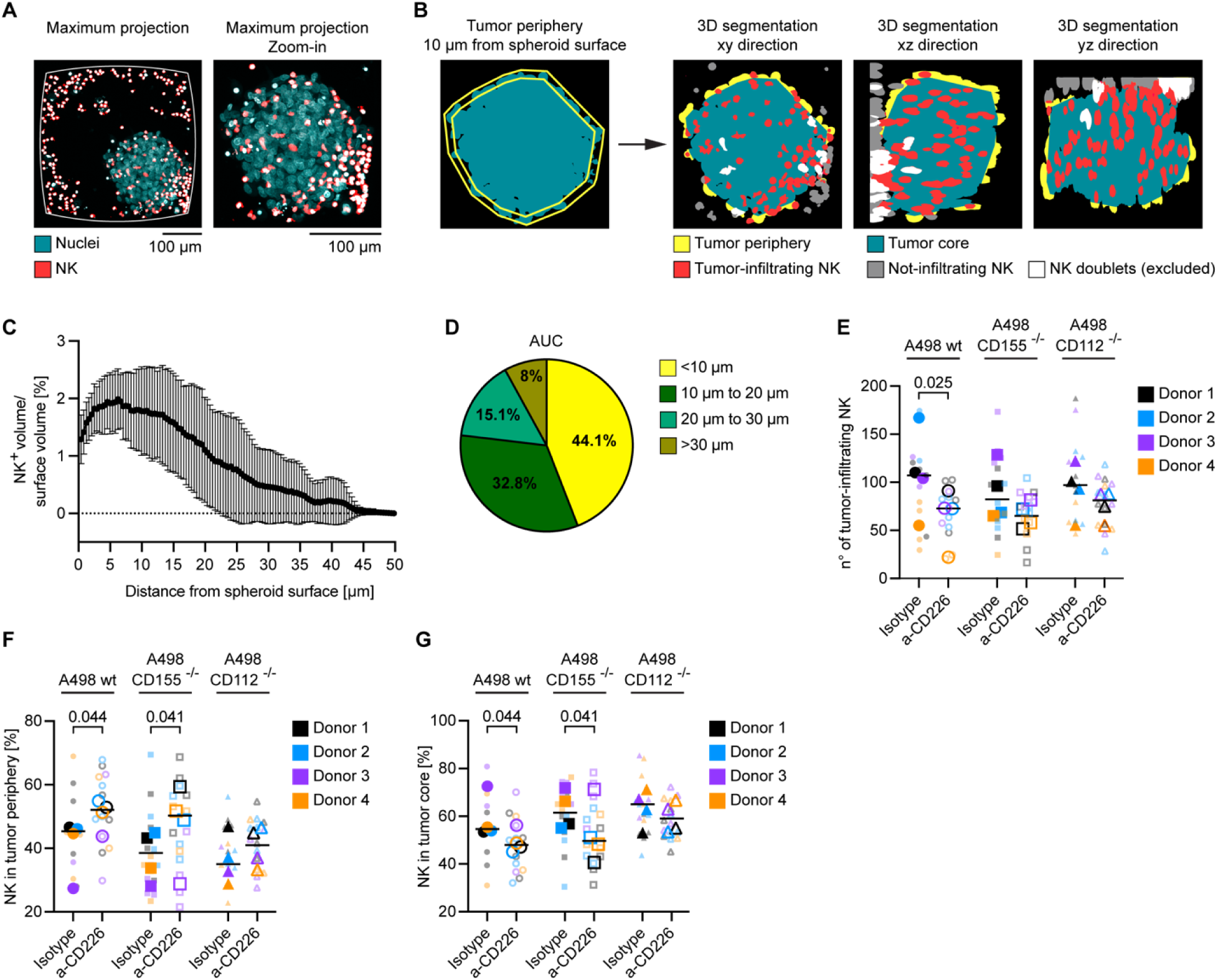
CD226 promotes NK cell infiltration. **A.** A498wt spheroids were maintained in culture for 48 hours before addition of IL-15-activated NK cells pre-stained with CellTrace Yellow (E:T seeding ratio = 1:1). After 5 hours, the co-cultures were fixed and permeabilized before proceeding with nuclear staining and spheroid clearing. 3D maximum projections of NK cells (in red) and nuclei (in cyan). **B.** A representative example of 3D spheroid segmentation in xy, xz and yz directions: spheroid periphery (<10 μm from the surface, in yellow) and spheroid core (>10 μm from the surface, teal). In red: NK cells. **C.** Profile of the volume occupied by NK cell signal (NK^+^ volume) normalized for the surface volume (y-axes) in relation to the distance from the spheroid surface (x-axes). **D.** Quantification of the Area Under the Curve (AUC) corresponding to each distance from the spheroid surface in relation to the Total AUC from C. **E.** Quantification of NK cell infiltration in the total spheroid area in A498 ^CD155wt^ ^CD112wt^ (circles), A498 ^CD155-/-CD112wt^ (squares) and A498 ^CD155wt^ ^CD112-/-^ (triangles) spheroids, in the presence (empty shapes) or in the absence (filled shapes) of CD226 blockade. The big symbols represent the mean values for 4 spheroids. Each color represents a different donor. Statistical method: two-way ANOVA followed by Tukey’s Post Hoc test (n=4). **F, G.** NK cell infiltration in the periphery (**F**) and in the core (G) of A498 ^CD155wt CD112wt^, A498 ^CD155wt CD112wt^ and A498 ^CD155wt CD112wt^ spheroids. A498 ^CD155wt CD112wt^ (circles), A498 CD155-/-CD112wt (squares) and A498 ^CD155wt^ ^CD112-/-^ (triangles) spheroids, in the presence (empty shapes) or in the absence (filled shapes) of CD226 blockade. The big symbols represent the mean values for 4 spheroids. Each color represents a different donor. Statistical method: two-way ANOVA followed by Tukey’s Post Hoc test (n=4).

Next, we evaluated the impact of CD226 on NK cell infiltration by incubating NK cells with A498 ^CD155wt CD112wt^, A498 ^CD155-/- CD112wt^ and A498 ^CD155wt CD112-/-^ spheroids with or without CD226 blocking. No differences in radii were observed between A498 ^CD155wt^ ^CD112wt^, A498 ^CD155-/-CD112wt^ and A498 ^CD155wt^ ^CD112-/-^at t0, indicating no impact of CD155 and CD112 in spheroid stability, growth and survival (Fig.S3Q). To estimate NK cell infiltration more precisely, we further develop the image analysis pipeline to provide the actual NK cell number in the spheroid. NK infiltration in the total spheroid volume significantly decreased in the presence of CD226 blockade in A498 ^CD155wt^ ^CD112wt^ (Fig.5E), indicating that CD226 promotes NK cell infiltration in solid tumors. If one of the CD226 ligands was not expressed by tumor cells, CD226 blockade was not significantly different from the isotype control (Fig.5E). Therefore, CD226 promotes NK cell adhesion and infiltration in tumor spheroids via both CD155 and CD112.

To better understand if CD155 and CD112 had distinct roles in promoting NK cell attachment to the tumor and subsequent infiltration, we quantified the number of NK cells in two separate spheroid regions (Fig. 5B): 1) tumor periphery, corresponding to the region from the spheroid surface and 10 μm inward (Fig.5B, in yellow); 2) core, defined as the inner region of the spheroid, deeper than 10 μm from the spheroid surface (Fig.5B, in teal).

CD226 blockade in A498 ^CD155wt^ ^CD112wt^ and A498 ^CD155-/-CD112wt^ spheroids caused an enrichment of NK cells in the tumor periphery, corresponding to a decrease of NK infiltration in the spheroid core (Fig.5F, G). The results indicate that CD226-CD112 ligation specifically promotes NK cell migration from the periphery to the tumor core. These findings are supported by our previous data showing that CD226 internalization was induced by CD155 but not by CD112, indicating the possibility of CD226 continuously engaging with CD112 towards the tumor core. The effect of CD226 blockade did not cause significant variations of NK cell distribution in the periphery and the core of in A498 ^CD155wt^ ^CD112-/-^ spheroids (Fig.5E, F), suggesting that CD155-CD226 ligation contributes equally to NK migration in both the periphery and the core of solid tumors. However, we cannot exclude that the absence of CD112R inhibition in A498 ^CD155wt^ ^CD112-/-^ spheroids could have enhanced NK activation and consequently increased the overall distribution of NK cells in the spheroid core. We were unable to verify this hypothesis in these settings due to the experimental complexity.

Overall, our data shows that CD226 promotes NK cell migration into the tumor tissues via CD155 and CD112 ligation, the latter being important for NK cell penetration into the spheroid core from the periphery. The findings also point towards the use of CD112R-CD112 blockade to simultaneously enhance NK cell killing activity and promote infiltration into the tumor core via CD226 and CD112 interaction.

## DISCUSSION

Strategies to target CD155 and CD112 have been proposed for two decades. However, a comprehensive understanding of CD155 and CD112 interaction with their immune cell receptors and the outcome of multiple integrating signals has been missing, delaying the introduction of CD155 and CD112-targeting therapies in clinics, and causing clinical trial failures. Expanding our insights of the CD155 and CD112 axes is crucial to identifying the most effective targets, to predict patient response and ultimately reduce treatment failure.

Combining the blocking of CD155 and CD112 receptors on NK cells with the use of renal carcinoma cell lines knock-out for either one or both ligands, we demonstrated that CD226 preferentially binds to CD155 rather than CD112, with TIGIT and KIR2DL5 overcoming CD226-mediated signaling. In case of KIR2DL5, this might be the result of direct ligand binding competition, while no evidence for this was found for TIGIT. Consistent with our conclusions, Zhu Y. et al. previously demonstrated that TIGIT had little effect on disrupting CD112R-CD112 interaction, while CD226 was a good inhibitor of the CD112-CD112R binding, binding indicating that CD112R and CD226 share a common binding site on CD112. In addition, Zhu Y. et al. used coated beads with individual Nectin-like proteins and stained for CD112R protein binding, finding that no PVR-like protein except CD112 was able to interact with the CD112R, supporting the notion that CD112 is the main ligand, if not the only one, that mediates the interaction of CD112R with DCs and tumor cells(*43*). While these studies are useful to understand binding sites and molecular interactions, they did not show how protein binding translates to NK cell responses. Our manuscript goes a step further by showing which interaction leads to degranulation responses in NK cells.

Despite the reported inhibitory nature of KIR2DL5 (*30*, *36*), its ligand was unknown for two decades and it was only recently identified as CD155 (*12*). We found that TIGIT tends to be expressed more in KIR2DL5^-^ NK cells compared to KIR2DL5^+^ NK cells, possibly balancing the inhibition input between the two populations.

Our results are in line with the “inhibitory receptor first” mechanism (*44*), i.e. that inhibitory receptors, such as TIGIT and KIR2DL5, prevail over the corresponding activating receptors, CD226 in this case. Accordingly, tumor growth and metastasis are reduced in CD155^-/-^ mice compared to CD155^wt^, likely due to CD155-mediated inhibition of T and NK cells (*33*). Simultaneous CD155 loss and CD112R receptor blockade enhanced the degranulation in TIGIT^+^ and KIR2DL5^+^ NK cells, indicating additive inhibitory activity induced by these three receptors. Similar increases in NK cell degranulation were observed using combinatorial blocking of TIGIT and CD112R, suggesting combinatorial treatment as a strategy to improve patient survival in CD155^+^ and CD112^+^ tumors.

While inhibitory receptors tend to bind to either CD155 or CD112, CD226 shows more plasticity in response to the environment, being able to switch between CD155 and CD112 depending on the ligand availability on tumor cells. This, together with the biological meaning of CD226 down-regulation induced by CD155 and not CD112, and its impact on NK cell-tumor cell contact duration and serial killing remains to be defined, and it will be the topic of future investigations. On the other hand, variations in CD155-expression influenced the levels of degranulation in TIGIT^+^ and KIR2DL5^+^ cells. High sensitivity to CD155 levels suggests that differences among individuals in terms of TIGIT, KIR2LD5 and intratumoral CD155 expression might lead to different outcomes and have a direct impact on the efficacy of anti-TIGIT therapy.

As the efficacy of checkpoint blockade depends on the availability of the target and the receptor, as well as on the ability of immune cells to infiltrate and perform cytotoxicity in a 3D solid tumor microenvironment, we tested the hierarchy of CD155 and CD112 receptor activation and its impact on TIGIT blockade in spheroid models of renal carcinoma.

TIGIT blockade improved the ability of NK cells to kill solid tumors, but it did not induce a complete spheroid eradication, leaving around 40% of viable tumor cells at the end of the treatment, mainly localized at the spheroid core. Interestingly, a reduction in CD155 expression accompanied the maturation of the spheroids, suggesting a role of cell-to-cell contact in CD155 modulation. Indeed, CD155 is involved in initiating the formation of adherent junctions via heterophilic interactions and is subsequently internalized through clathrin-dependent endocytosis(*5*, *45*). Despite a consistent presence of TIGIT^+^ NK cells infiltrating the spheroids, TIGIT expression significantly decreased in this subset.

TIGIT modulation was also observed after the treatment with lenalidomide and axitinib, causing loss of efficacy in combinatorial treatment with TIGIT blockade. In line with our observations, it has been shown that lenalidomide exposure decreases the threshold of NK cell activation (*46*) and improves ADCC against CD155^+^ and MICA^+^ tumor cell lines (*47*). Our data shows that low CD155 availability combined with decreased expression of TIGIT induced by the tumor microenvironment and previous drug exposure might reduce the impact of TIGIT-mediate immune cell inhibition in solid tumors, potentially affecting TIGIT blockade long-term efficacy and causing treatment failure.

On the other hand, CD112 expression in tumor spheroids was stable, and the combination of NK therapy with CD112R blockade induced rapid and durable responses. Together this suggests that TIGIT and CD155 expression in the tumor microenvironment both before and after treatment with other therapies should be carefully taken into consideration before selecting patients for anti-TIGIT therapy, and CD112R blockade could be considered as a better option due to less CD112 modulation by the solid tumor architecture.

Immune-cell infiltration into tumor tissue is a critical requirement for successful tumor therapy. Previous reports described CD226 involvement in regulating trans-endothelial migration of monocytes via CD155 interaction (*48*), but no direct evidence of CD226 contribution to NK cell tumor infiltration has been available to date. Here we provide a comprehensive overview of CD226-mediated regulation of NK cell migration in tumor tissues, by showing that CD226 promotes NK cell adhesion and infiltration in tumor spheroids via both CD155 and CD112, with CD226-CD112 ligation specifically driving NK cells from the periphery to the tumor core. As CD155 expression decreased with cell-to-cell contact-maturation in the tumor spheroids, one could speculate that this drives the switch from CD155 to CD112 binding. In addition, CD226 internalization induced by CD155 but not by CD112 might favor continuous engagement with CD112 towards the tumor core. Further investigation is needed to elucidate the biological meaning of CD226 plasticity observed in both 2D and 3D settings.

Overall, our findings point towards the use of CD112R-CD112 blockade to simultaneously enhance NK cell killing activity and promote infiltration into the tumor core via CD226 and CD112 interaction. On the other hand, TIGIT blockade might be more effective in the context of tumor metastasis, where loss of cell contact inhibition promotes CD155 overexpression. In line with that, CD155 immunostaining is usually higher in metastatic tumors compared to primary tumors (*14*, *49*).

In this study, we provide deep functional understanding of CD155 and CD112 receptors within the solid tumor microenvironment and their effects on cellular immunotherapy and immune-cell infiltration. Our findings may help inform how, when, and in whom these therapies should be used, either alone or in combination with other agents.

## MATERIALS AND METHODS

### NK cell isolation and cell culture

NK cells were obtained from buffy coats by negative selection using EasySep Direct Human NK cell isolation kit (Stemcell Technology) according to manufacturer’s instructions and maintained in RPMI 1640 with Lglutamine (Sigma-Aldrich) supplemented with 10% fetal bovine serum (FBS, Sigma-Aldrich), 1x MEM Non-Essential Amino Acid Solution (Sigma-Aldrich) and 10 mM Hepes (Sigma-Aldrich) and 10 ng/mL of IL-15 (R&D System). To study the effect of axitinib and lenalidomide on TIGIT expression, NK cells were incubated for 48 hours with 10 ng/mL of IL-15 and 5 µM of axitinib (Sigma-Aldrich) or 1 µM of lenalidomide (Sigma-Aldrich).

A498 cells (ATCC, RRID: CVCL_1056) were maintained in RPMI 1640 with L-glutamine (Sigma-Aldrich) supplemented with 10% FBS, 1x MEM Non-Essential Amino Acid Solution and 10 mM Hepes at 37 °C in 5% CO_2_ and passaged before they reached confluency. A498 cell line has been validated using PCR-based single-locus technology.

### Spheroid formation and culture

The ultrasound-induced spheroid formation in the multichambered microwell chip has been previously described (*38*, *50*). Miniaturized agarose-induced 3D cultures: agarose micro-well hydrogels obtained with spheroid micro-mold (#12–256-Small, MicroTissues, Inc., well diameter = 400 μm) were equilibrated in complete culture medium. A498 renal cell carcinoma cells were seeded at 1×10^6^ cells/mL (100 μl per gel) and allowed to settle in the microwells for 30 min, then 2 mL of complete medium were added. Ultrasound-and agarose-induced spheroids were maintained at 37 °C, 5% CO_2_ before being harvested for further analysis.

### Spheroid collection and treatment for flow cytometry analysis and functional assays

Ultrasound-induced and agarose-induced spheroids were harvested, washed three times with PBS (Sigma-Aldrich) before performing enzymatic dissociation for 45 minutes at room temperature using Accumax (Stemcell Technologies). After Accumax treatment, cells were washed twice with complete medium and resuspended in staining buffer for immunostaining or used directly in CD107a degranulation assay as target cells. The same procedure, including accumax treatment, was performed on A498 cells cultured as monolayers, on NK cells cultured alone and in co-culture with A498 spheroids.

### Immunostaining and CD107a degranulation assay

Immunostaining for flow cytometry and confocal microscopy was performed as previously described(*51*). CD107a assay was performed as following: overnight IL-15 activated NK cells and A498 cells were harvested, counted, re-suspended in complete medium and transferred into a 96 V-bottom well plate (1×10^5^ cell/well, E:T ratio = 1:1). To assess spontaneous NK-cell degranulation, control samples without A498 target cells were included. Spontaneous degranulation was subtracted from all measurements, and the data presented in the paper are normalized values. For antibody blocking, a 30-minutes pre-incubation step with the neutralizing antibodies was performed before co-culturing effector and target cells. Cells were incubated at 37°C, 5% CO_2_ in the dark for 2 hours in the presence of anti-CD107a antibody, before proceeding to immunostaining and flow cytometry analysis. Data were acquired by BD FACSCanto IVD 10, standard 3-lasers, 10 colors (BD Biosciences) and analyzed with the FlowJo Software v10 (FlowJO, LLC, RRID:SCR_008520). The following gating strategy was applied to identify degranulating NK cells: after separating NK cells from tumor cells by size and granularity (forward scatter vs side scatter), the following gates were created: singlets (forward scatter-A vs forward scatter-H): live cells: CD45^+^: CD3^-^: CD107a^+^. When indicated, CD107a^+^ cells were analyzed in CD45^+^: CD3^-^: TIGIT^+/-^, CD96^+/-^ and KIR2DL5^+/-^ gates.

### Live cell imaging of spheroid viability and analysis of NK cell infiltration and synapse

Live cell imaging of spheroid cytotoxicity was performed as previously described (*38*). Where indicated, blocking antibodies and drug treatments were added at the beginning of the assay (the list of treatments and concentrations is available in Supplementary Materials). For NK cell infiltration assays and immune synapse formation, NK cells were pre-stained with CellTrace Yellow (CTY, ThermoFischer Scientific) following the manufactureŕs instructions. The multichambered microwell chip was exposed to a short (1 hour) round of USW to focus the NK cells onto the tumor spheroids, before being transferred to a widefield microscope equipped with environmental control (37°C, 5% CO_2_ and humidity) for live cell timelapse imaging acquiring image frames every hour for 5 hours. At the end of the timelapse, spheroids were fixed with 4% (w/v) formaldehyde solution for 15 minutes and washed with PBS. Incubation with increasing concentration of methanol (50% in PBS, 80% in deionized water) and spheroid rehydration were performed before proceeding with DAPI staining (ThermoFisher Scientific, 2 μg/mL). Spheroid clearing was performed prior to imaging as previously described (*50*). The analysis of NK infiltration is described in Supplementary Material. To evaluate the recruitment of CD112 at the immune synapse, A498 cells were cultured overnight in 8-well chamber slides (Ibidi) to promote adhesion to the glass. NK cells were added to the culture for 30 minutes before proceeding to fixation, permeabilization and immunostaining as previously described. The detailed analysis of CD112 recruitment at the region of NK-tumor cell contact is available in Supplementary Material.

### Microscopy

Images of fixed samples were acquired by confocal microscopy (Zeiss LSM 880) using a 10x/0.45 Plan-Apochromat Air objective, a 20×/0.8 Plan-Apochromat Air objective or a 63×/1.40 Plan-Apochromat Oil immersion objective (Zeiss). Time-lapse live cell imaging was performed in a widefield Zeiss Axio Observer Z1 7 microscope equipped with an incubation chamber with environmental control (37°C, 5% CO2, and humidity), a Fluar 10×/0.50 Air objective and a Hamamatsu ORCA-flash 4.0 camera.

### Statistical analysis

Statistical analysis and curve fitting was performed using GraphPad Prism (GraphPad Software, La Jolla, CA, USA, RRID:SCR_002798). Comparison between two or multiple conditions was performed using unpaired t-test, One-way or Two-way ANOVA followed by Tukey’s Post Hoc test as indicated in the figure legends. Data groups were considered not significantly (n.s.) different when P > 0.05, and only significantly different annotations are shown in the figures. Normality tests were performed before proceeding to ANOVA and t-tests.

## LIMITATIONS OF THE STUDY

This study has two main limitations. This study has two main limitations. First, we used tumor spheroids as a model of solid tumors. Although 3D tumor spheroids are well-established *in vitro* platforms that reproduce key features of tumor biology, including tissue architecture, cell-cell and cell-ECM interactions, and drug diffusion, they do not fully capture the complexity of the tumor microenvironment. Nevertheless, these models allowed us to recapitulate the downregulation of CD155 observed in primary tumors compared with metastases and to investigate NK cell infiltration. Further advances in model systems are needed to better reflect the complexity of in vivo tumors. Second, anti-TIGIT and anti-CD112R antibodies were used during the degranulation assays to understand the effect of TIGIT and CD112R blockade. The presence of blocking antibodies for TIGIT and CD112R precluded the possibility to distinguish TIGIT+/- and CD112R +/- populations due to antibody occupancy. Different antibodies were tested to find alternatives to label TIGIT and CD112R simultaneously with two different antibodies, however total or partial occupancy was always observed, not allowing this type of experiment. Despite these limitations, we believe that the manuscript sheds new light on the dynamic interplay between CD155, CD112 and their NK cells receptors and provides a more comprehensive understanding of how the PVR/nectin family of proteins could be targeted in solid tumor immunotherapy.

## ACKNOLEDGEMENTS

We thank The Knut and Alice Wallenberg foundation, The Swedish research council, The Swedish foundation for strategic research, The Swedish childhood cancer foundation, The Swedish cancer foundation, The Cancer Research Funds of Radiumhemmet for financial support, Madoka Takai group for providing the coating polymer used in the multichambered microwell chip, and Benedict Chambers for useful discussions and for providing assistance during the manuscript preparation.

## Author contributions

Conceptualization: VC, BÖ

Methodology: VC, KO, HZ: NS; MW, BÖ

Investigation: VC, KO, AKW, JF, DT, GT, BH, HVO

Software: VC, KO, HZ, HVO

Formal analysis: VC

Resources: PAS, AL, MW, BÖ

Visualization: VC, BÖ

Funding acquisition: BÖ

Project administration: VC, BÖ

Supervision: AL, MW, BÖ

Writing – original draft: VC, BÖ

Writing – review & editing: VC, KO, HZ, NS, PAS, HVO, AL, MW, BÖ

## Data and materials availability

Correspondence and material requests should be addressed to Björn Önfelt and Valentina Carannante

## Additional information

### Financial support

The Knut and Alice Wallenberg foundation Grant 2018.0106

The Swedish research council Grant 2019-04925

The Swedish foundation for strategic research Grant SBE13-0092

The Swedish childhood cancer foundation Grant MT2019-0022

The Swedish cancer foundation Grant 19 0540 Pj and 21 1524 Pj

The Cancer Research Funds of Radiumhemmet Grant #211253

## SUPPLEMENTARY MATERIALS

### Materials and Methods

List of antibodies, fluorescent dyes and drugs

Generation of stable A498-RFPcells and CD155-KO A498 cells

Transient small interference RNA (siRNA)-mediated CD155 knockdown

NK isolation from KIR2DL5^+^ donors

Analysis script for NK infiltration

Analysis of CD112 recruitment at the region of NK-tumor cell contact

### Supplementary figures

Fig. S1. Characterization of CD155 and CD112 axes il IL-15 activated NK cells

Fig. S2. Characterization of NK and tumor responses to CD155 silencing on A498 ^CD155wt^ ^CD112wt^ renal carcinoma cells

Fig. S3. Characterization of CD155 and CD112 impact on NK cytotoxic activity and infiltration is solid tumor models with combinatorial treatments

### Supplementary Tables

Table S1. Percentages of degranulating NK cells gated on TIGIT+ and TIGIT-co-cultured with A498wt cells during CD155-silencing and blocking of CD226 (mean ± standard deviation)

Table S2. Percentages of total degranulating NK cells co-cultured with A498wt cells during CD155-silencing and blocking of CD226 (mean ± standard deviation)

## Notes

### Competing Interest Statement

The authors have declared no competing interest.

