## Supplementary Materials and Figures for "The solid tumor microenvironment changes the hierarchy of CD155 and CD112 receptors, shaping checkpoint blockade outcome"

**Supplementary Materials and methods**

**List of antibodies, fluorescent dyes and drugs**

The following antibodies were used for flow cytometry: FITC/Pe.Cy5/V500-conjugated anti-CD45 (HI30, RRID:AB_395874, RRID:AB_395876, RRID:AB_1937324), FITC-conjugated anti-CD95 (clone: DX2, RRID:AB_400038), PE-conjugated anti-TRAIL-R1 (clone: S35-934, AB_2738648), PE-conjugated anti-TRAIL-R2 (clone: YM366, AB_2732871), PE-conjugated MICA/B (clone: 6D4, AB_397077), Pe-Cy7/APC-H7/BV605-conjugated anti-CD3 (clone: SK7, RRID:AB_400215, RRID:AB_1645730, RRID:AB_2714001), APC-H7/BV605-conjugated anti-CD16 (clone: 3G8, RRID:AB_1727432, RRID:AB_2740165), APC-conjugated anti-PD-L1 (clone: 29E.2A3, RRID:AB_2916833), Alexa Fluor 647/BV421-conjugated anti-CD107a (H4A3, RRID:AB_2737684), BV421-conjugated CD56 (clone: BI59), BV421-conjugated anti-CD226 (clone: DX11, RRID:AB_2740826), BV421-conjugated anti-ICAM-1 (clone: HA58, RRID:AB_2738578), BV421-conjugated anti-EGFR (clone: EGFR.1, RRID:AB_2739632), BV605-conjugated anti-HLA-A,B,C (clone: G46-2.6, RRID:AB_2740137) and BV605-conjugated anti-TRAIL-R3 (clone: B-D44, RRID:AB_2742774) were purchased from purchased from BD Bioscience.PE-Cy7-conjugated anti-CD96 (clone REA196, RRID:AB_2751832), APC-conjugated anti-KIR2DL5 (clone: UP-R1, RRID:AB_2660360) and APC-conjugated anti-Nectin-4 (clone: REA967, RRID:AB_2727314) were purchases from Miltenyi Biotech. PE/Alexa Fluor 647/BV605-conjugated-anti-TIGIT (clone: VSTM3, RRID:AB_10895760, RRID:AB_2715972, RRID:AB_2632926), Alexa-fluor 488-conjugated MICA/B (clone: 6D4, RRID:AB_2143627), PE-conjugated anti-CD56 (clone: HCD56, RRID:AB_604093), PE-conjugated anti-Nectin-2 (clone: TX31, RRID:AB_2269088) and APC-conjugated CD155 (clone: SKII4, RRID:AB_2565815) were purchased from Biolegend. Purified anti-CD45 (clone EP322Y, RRID:AB_726545) was purchased from Abcam. Rabbit polyclonal anti-Nectin-1 (cat. numb. HPA026846, RRID:AB_1846226) and rabbit polyclonal anti-Nectin-3 (cat. numb. HPA011038, RRID:AB_1079719) were purchased from Atlas Antibodies. The following antibodies were used for confocal microscopy: polyclonal goat anti-human CD112 (Clone: AF2229, R&D Systems, RRID: AB_2269089) and donkey anti-goat Alexa Fluor 647 (Clone ab150135, Abcam, RRIS: AB_2687955). The following blocking antibody were used in the degranulation assays and image-based NK killing assays: mouse anti-human CD155 (clone D171, Thermo Fischer Scientific), mouse anti-human TIGIT (Clone: MBSA43; eBioscience, ThermoFisher Scientific, RRID:AB_10978147), mouse anti-human CD226 (Clone: DX11, BD Bioscience, RRID:AB_397328), mouse anti-human CD96 (clone: NK92.39, Abcam, RRID:AB_1658502), mouse anti-Nectin-2 (Clone: TX31, Biolegend, RRID:AB_2174164) and mouse igG1 k isotype control (clone: P3.6.2.8.1, eBioscience, ThermoFisher Scientific, RRID:AB_470111) at final concentration of 10 µg/mL. The following dyes have been used: DAPI (final concentration: 2 µM, ThermoFisher Scientific), and CellTrace yellow (final concentration: 2.5 µM, ThermoFisher Scientific), TMRM (final concentration: 200 nM; Invitrogen, ThermoFisher Scientific) and Oregon Green 488 Phalloidin (final concentration: 60 nM, Sigma-Aldrich). The following drug have been added at the beginning of NK cell cytotoxicity live imaging assay: 5 µM of axitinib (Sigma-Aldrich), 1 µM of lenalidomide (Sigma-Aldrich).

**Generation of stable A498-RFPcells and CD155-KO A498 cells**

The stable A498 cell line expressing RFP (A498-RFP) was obtained using a pLenti-CMV-MCS-RFP-SV-puro vector as previously described(*1*). A498-RFP cells were cultured in complete medium with 2 µg/mL puromycin.

To generate A498 cells with a knockout of CD155, gRNAs were designed to bind within either exon 1 (GGGGGTCACTCACCGGTTCC) or exon 2 (CTATTCGGAGTCCAAACGGC) of the gene using the CRISPOR algorithm(*2*). The gRNAs were cloned into the lentiviral expression vector lentiCRISPRv2 using BsmBI (NEB) insertion sites(*3*). Plasmids were verified by sequencing. Lentivirus was produced as previously described(*4*). Briefly, $1x10^6$ HEK293FT cells were plated into a poly-D-lysine-coated 60-mm dish (Corning) The following day, the cells were transfected with 4.8 µg of the cloned lentiCRISPRv2 containing the gRNAs, 2.4 µg of pMDLg/pRRE (#12251 Addgene, RRID:Addgene_12251), 1.6µg of pRSV-REV (#12553 Addgene, RRID:Addgene_12253), and 0.8 µg of pCMV-VSV-G (#8454 Addgene, RRID:Addgene_8454) using calcium phosphate transfection kit (Sigma-Aldrich) in the presence of 25µM chloroquine (Sigma-Aldrich). The medium was changed 14 hours post-transfection, and the virus particles were collected after an additional 24 hours, by filtering the supernatant through a 0.45 µm filter and stored at -80°C. 8000 A498 cells suspended in 2mL of RPMI medium supplemented with 10% FBS per condition were plated and allowed to adhere to a 6-well plate (Corning). 16 hours post-plating, they were transduced with 100 µl of virus-containing supernatant for 6 hours in 8 µg/mL protamine sulfate (Sigma Aldrich). After transduction, cells were cultured for 7 days and then stained using anti-CD155 antibody (clone SKII.4, Biolegend). The cells were sorted using fluorescence-activated cell sorting (FACS) with BDARIA Fusion (BD Biosciences). Sorted cells were cultured additional 7 days before using them in experiments.

**Transient small interference RNA (siRNA)-mediated CD155 knockdown**

Here we used FlexiTube GeneSolution siRNA for CD155 (Qiagen, cat.numb. 1027416) containing four siRNA (Hs_CD155_1; Hs_CD155_5; Hs_CD155_7; Hs_CD155_8) targeting the following CD155 mRNA sequences:

Hs_CD155_1: CAACACAACTTTAATCTGCAA
Hs_CD155_5: CAGGCTATAATTGGAGCACGA
Hs_CD155_7: TCCTGTGGACAAACCAATCAA
Hs_CD155_8: CGGCAAGAATGTGACCTGCAA

Negative control siRNA (Quiagen, cat.numb.1027310) was used as control. 100 000 A498 cells were cultured in T25 flasks in complete culture medium and left to adhere to the flask surface overnight (day 0). On day 1 (96 hours incubation), day 2 (72 hours), day 3 (48 hours) and day 4 (24 hours), adherent A498 cells were incubated with 10 nM solution of FlexiTube GeneSolution siRNA for CD155 or 10 nM solution of negative control siRNA using OptiMEM reduced serum medium (Gibco, ThermoFisher Scientific) and Lipofectamin RNAiMAX

(Invitrogen, ThermoFisher Scientific) according to manufacturer's instructions. After 8 hours, A498 cells were washed twice and maintained in complete culture medium. On day 5, cells were detached with Accumax (Stemcell Technologies, 5-minute incubation at room temperature) and used for flow cytometry analysis of NK cell ligand expression and CD107a release assays.

##### NK isolation from KIR2DL5$^+$ donors

To obtain NK cells from KIR2DL5$^+$ donors we performed a two-step separation: first, peripheral blood mononuclear cells (PBMCs) were isolated by Ficoll–Hypaque gradient separation (GE-Healthcare), followed by flow cytometry screen for KIR2LD5 expression. When KIR2DL5 expression was confirmed, NK cell isolation from PBMCs was performed by negative selection using NK cell Isolation Kit (Miltenyi Biotec GmbH, GE). A two-step separation was also performed for KIR2DL5$^-$ donors used as control, to avoid artifacts due to technical variation.

##### Analysis script for NK infiltration

The NK cell infiltration into A498 spheroids was quantified by image analysis of confocal z-stacks (z-step size: 1 µm). The tumor spheroid was defined as a binary mask of nucleus segmentation followed by dilation processing, and NK cells as a label image (0=background, 1=single NK, 2=NK cluster). Volumes were resampled to a quasi-isotropic grid using an axial scale corrected for refractive-index mismatch. The spheroid mask was regularized and converted to a convex interior to enable robust "contour-to-center" distance statistics. Three biologically interpretable zones were generated around the spheroid boundary (outside, surface ring, inner core) using a user-set physical thickness (e.g., 10 µm). Inside the convex interior, a distance-from-surface map defined 1-voxel shells, from which we computed per-depth NK distribution and equal-volume radial partitions to remove surface area bias. Object-level summaries (counts, volumes, circularity, estimated cell numbers) were derived per zone. The MATLAB source codes are available in GitHub (link: [https://github.com/Hanqing-Zhang-KTH/Single-Spheroid-Infiltration-Analysis](https://github.com/Hanqing-Zhang-KTH/Single-Spheroid-Infiltration-Analysis)) and can be used to generate results and overlays for downstream analysis. The key quantitative steps in the analysis script are described here:

**Data and segmentation.**

Voxel sizes ($\Delta x$, $\Delta y$, $\Delta z$) are in µm. Spheroid mask, NK label (0=background, 1=single, 2=cluster), together with the 3D cell intensity stack (Channel 1= Tumor spheroid signal, Channel 2= NK cells signals, Channel 3 = brightfield) are used as input for the analysis script.

**Axial scaling and RI correction (quasi-isotropic resampling).**

Stacks are resampled so that the analysis grid is approximately isotropic; labels with nearest-neighbour, intensities with linear interpolation. The axial scale factor is:

$$s_z = \frac{\Delta z}{\max(\Delta x, \Delta y)} \cdot \alpha_{\mathrm{RI}}$$

**Spheroid regularization and convex interior.**

On a coarse grid, we morphologically smooth $S_0$, keep the largest component, and compute its 3D convex hull S, re-embedded at full resolution and used as the interior domain.

**Fixed-thickness zones (±r around the surface).**

Let r be the desired thickness in voxels (r ≈ round (check_range_um / Δx)) and $B_r$ a spherical structuring element. Zones are defined by:

| | |
|---|---|
| (S ⊕ B_r) \ S | *outside zone* |
| S \ (S ⊖ B_r) | *surface zone* |
| S ⊖ B_r | *inner zone* |

**Distance-from-surface map and shells.**

Let ∂S be the 3D surface voxels. Define the Euclidean distance inside S and quantize to 1-voxel shells:

$$D(\mathbf{x}) = \mathrm{dist}(\mathbf{x}, \partial S),\ \ \mathbf{x} \in S$$

$$D(\mathbf{x}) = 0,\ \ \mathbf{x} \notin S$$

$$L(\mathbf{x}) = \mathrm{round}(D(\mathbf{x}))$$

**Radial (contour-to-center) statistics.**

For each shell d=1,…,$d_{max}$ we compute shell size, NK-single voxels, NK-cluster voxels, empty voxels, and all NK occupancy ratio:

$$N_{\mathrm{shell}}(d) = \#\{\mathbf{x} : L(\mathbf{x}) = d\}$$

$$N_{\mathrm{NK1}}(d) = \#\{\mathbf{x} : L(\mathbf{x}) = d \wedge \mathrm{label} = 1\}$$

$$N_{\mathrm{NK2}}(d) = \#\{\mathbf{x} : L(\mathbf{x}) = d \wedge \mathrm{label} = 2\}$$

$$N_{\mathrm{empty}}(d) = N_{\mathrm{shell}}(d) - N_{\mathrm{NK1}}(d) - N_{\mathrm{NK2}}(d)$$

$$f_{\mathrm{NK}}(d) = \frac{N_{\mathrm{NK1}}(d) + N_{\mathrm{NK2}}(d)}{N_{\mathrm{shell}}(d)}$$

**Equi-volume partitions.**

Let C(*d*)=∑{k≤*d*} $N_{shell}$(k) and $V_{tot}$=C($d_{max}$). For K bins, boundary indices $b_i$ satisfy C($b_i$)≈i·$V_{tot}$/K. Within each bin, recompute NK counts and occupancy $f_{NK}$.

$$C(d) = \sum_{k \le d} N_{\mathrm{shell}}(k), \quad V_{\mathrm{tot}} = C(d_{\mathrm{max}})$$

**Object-level zone summaries.**

For each NK object o (connected component) with voxel set $\Omega_o$, we compute which zone each object belongs to as the following:

$$V_o = |\Omega_o|, \quad \hat{n}_o = \max(1, \lfloor V_o / V_{\mathrm{single}} \rfloor)$$

$$\mathrm{zone}(o) = \arg \max_{z \in \{\mathrm{outside, surface, inner}\}} |\Omega_o \cap \mathrm{zone}_z|$$

**Analysis results details:**

Direct results from the analysis script include:

• Distributions (per-shell): distance (μm), shell voxels, empty, NK single, NK cluster, NK all. Interpretation: radial profiles from surface toward center at single-voxel resolution.
• EquiVolumeDistribution: same rows using equal-volume bins; interpretation: This NK distribution vs normed bin step profiles are robust to spheroid size/shape and can be used to compare averaged samples.
• ALL_Cells: per-zone object counts, summed volumes, volume % per zone, mean object volume, class totals, and metadata.
• ALL_Cells_IsoVoxelDetails: per-object volume, circularity, centroid, class, estimated cell number, and assigned zone.
• Overlaying PNG: max projection showing convex outline, surface bands, and optional spheroid/NK overlays.

**Post-analysis**

1) NK-all occupancy vs Constant depth step: For NK distributions, we compute the NK occupancy ratio $f_{NK}$ and plot it versus distance from the surface. Averaging with window size of 3 distance steps is used to smooth the data.
2) 10-μm zone summaries: From tumor spheroid and NK data, bar plots per zone (outside/surface/inner) of NK counts, summed volumes, volume %, and mean object volume are collected.

#### Analysis of CD112 recruitment at the region of NK-tumor cell contact

To evaluate CD112 recruitment at the region of NK-tumor contact, z-stack images of NK-A498 cell conjugates were acquired at 0.5 μm intervals. The number of optical sections was determined by setting the upper and lower z-axis limits to include the entire conjugate. In ImageJ, the region of contact was defined as the region of F-actin accumulation (identified by phalloidin staining) at the interface between conjugates. Specificallt, z-stacks images corresponding to 3 μm volume centered on the plane showing the highest F-actin intensity were selected and the sum intensity projection generated to identify the regiond of interest (ROI) using the 'Analyze Particles' function in ImageJ. Mean fluorescence intensity in the CD112 channel was then measured from the average-intensity projection within the ROI. The data were normalized to the background fluorescence by subtractin the mean fluorescence intensity from the CD112 channel in the control condition (NK incubated with A498 $^{CD155wt\,CD112-/-}$ cells).

### Supplementary figures

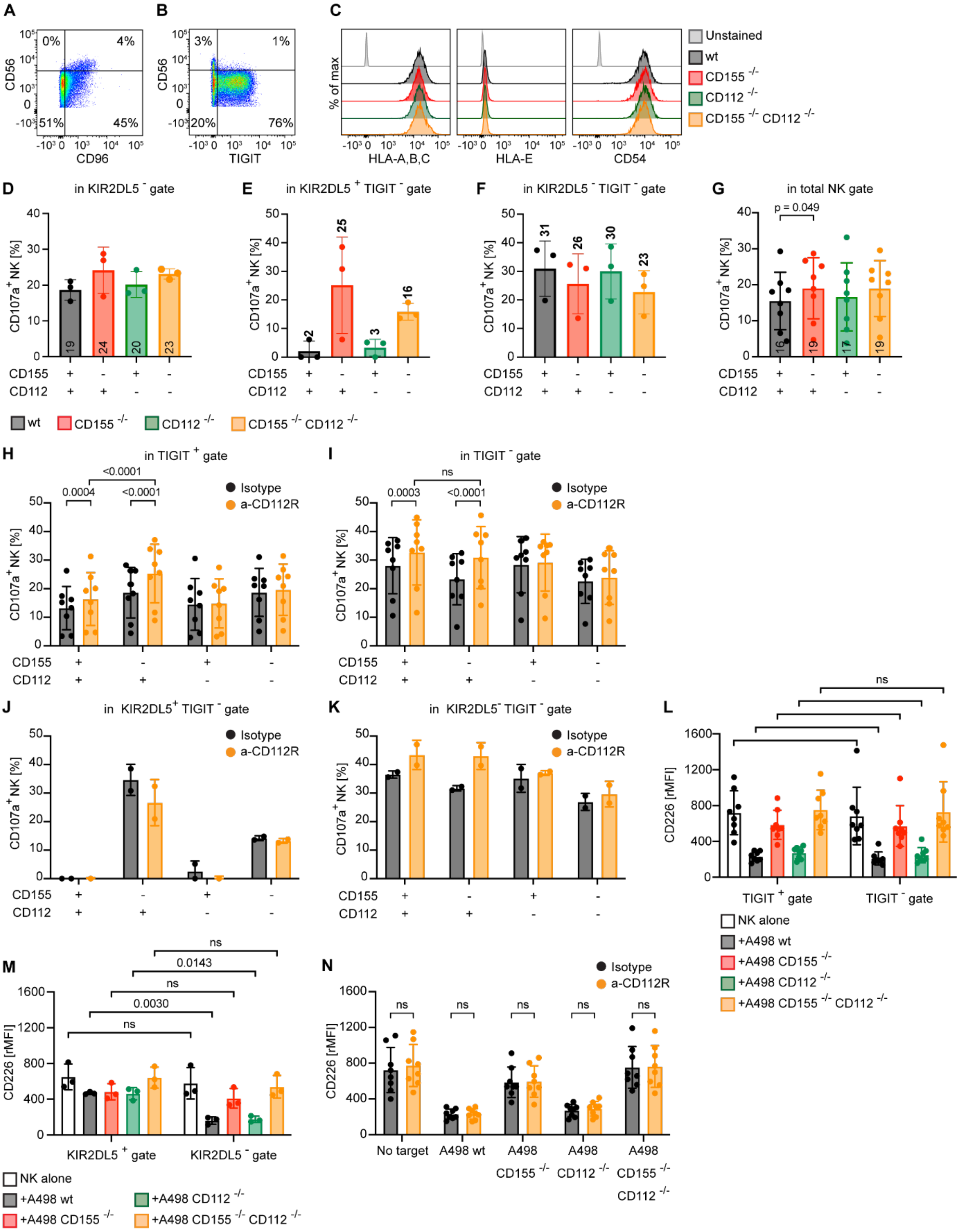


**Fig. S1. Characterization of CD155 and CD112 axes in IL-15 activated NK cells**

**A, B.** Flow cytometry dot-plots showing the expression of CD96 (**A**) and TIGIT (**B**) in relation to $CD56^{Bright}$ and $CD56^{Dim}$ NK cell populations after overnight activation with IL-15 (10 ng/mL). Percentage of positive cells are indicated within the corresponding gates. **C.** HLA Class I, HLA-E and CD54 surface expression on

A498 $^{CD155wt\ CD112wt}$ (black histogram), A498$^{CD155-/-\ CD112wt}$ (red histogram), A498$^{CD155wt\ CD112-/-}$ (green histogram) and A498$^{CD155\ -/-\ CD112-/-}$ (orange histogram). Unstained control is shown in light grey. **D-F.** Percentage of degranulating NK cells in response to A498 $^{CD155wt\ CD112wt}$ (black bar), A498$^{CD155-/-\ CD112wt}$ (red bar), A498$^{CD155wt\ CD112-/-}$ (green bar) and A498$^{CD155\ -/-\ CD112-/-}$ (orange bar) in KIR2DL5$^-$ (**D**) KIR2DL5$^+$ TIGIT$^-$ (**E**) KIR2DL5$^-$ TIGIT$^-$ (**F**) gates (n=3). **G.** Percentage of degranulating NK cells in response to A498 $^{CD155wt\ CD112wt}$ (black bar), A498$^{CD155-/-\ CD112wt}$ (red bar), A498$^{CD155wt\ CD112-/-}$ (green bar) and A498$^{CD155\ -/-\ CD112-/-}$ (orange bar) in NK gate (n=8). **H-K.** Effect of CD112R blockade (orange bars) on NK cell degranulation in TIGIT$^+$ (**H**), TIGIT$^-$ (**I**), KIR2DL5$^+$ TIGIT$^-$ (**J**) KIR2DL5$^-$ TIGIT$^-$ (K) gates after exposure to A498 $^{CD15wt\ CD112wt}$ (+,+), A498$^{CD155-/-\ CD112wt}$ (-,+), A498$^{CD155wt\ CD112-/-}$ (+,-) and A498$^{CD155\ -/-\ CD112-/-}$ (-,-) as indicated on the x-axis (n=8 in H, I; n=2 in J, K). Statistical method in H-I: two-way ANOVA followed by Tukey's Post Hoc test. **L.** CD226 surface expression on NK cells after exposure to A498wt (black bar), A498$^{CD155-/-\ CD112wt}$ (red bar), A498$^{CD155wt\ CD112-/-}$ (green bar) and A498$^{CD155\ -/-\ CD112-/-}$ (orange bar) compared to no target control (white bar) in TIGIT$^+$ gate (left panel) and TIGIT$^-$ gate (right panel) (n=8). **M.** CD226 surface expression on NK cells after exposure to A498wt (black bar), A498$^{CD155-/-\ CD112wt}$ (red bar), A498$^{CD155wt\ CD112-/-}$ (green bar) and A498$^{CD155\ -/-\ CD112-/-}$ (orange bar) compared to no target control (white bar) in KIR2DL5$^+$ gate (left panel) and KIR2DL5$^-$ gate (right panel) (n=3). **N.** CD226 surface expression on NK cells alone or after exposure to A498 $^{CD15wt\ CD112wt}$, A498$^{CD155-/-\ CD112wt}$, A498$^{CD155wt\ CD112-/-}$ and A498$^{CD155\ -/-\ CD112-/-}$ in the presence (orange bars) or in the absence (black bars) of anti-CD112R (n=8). Statistical method: two-way ANOVA followed by Tukey's Post Hoc test.

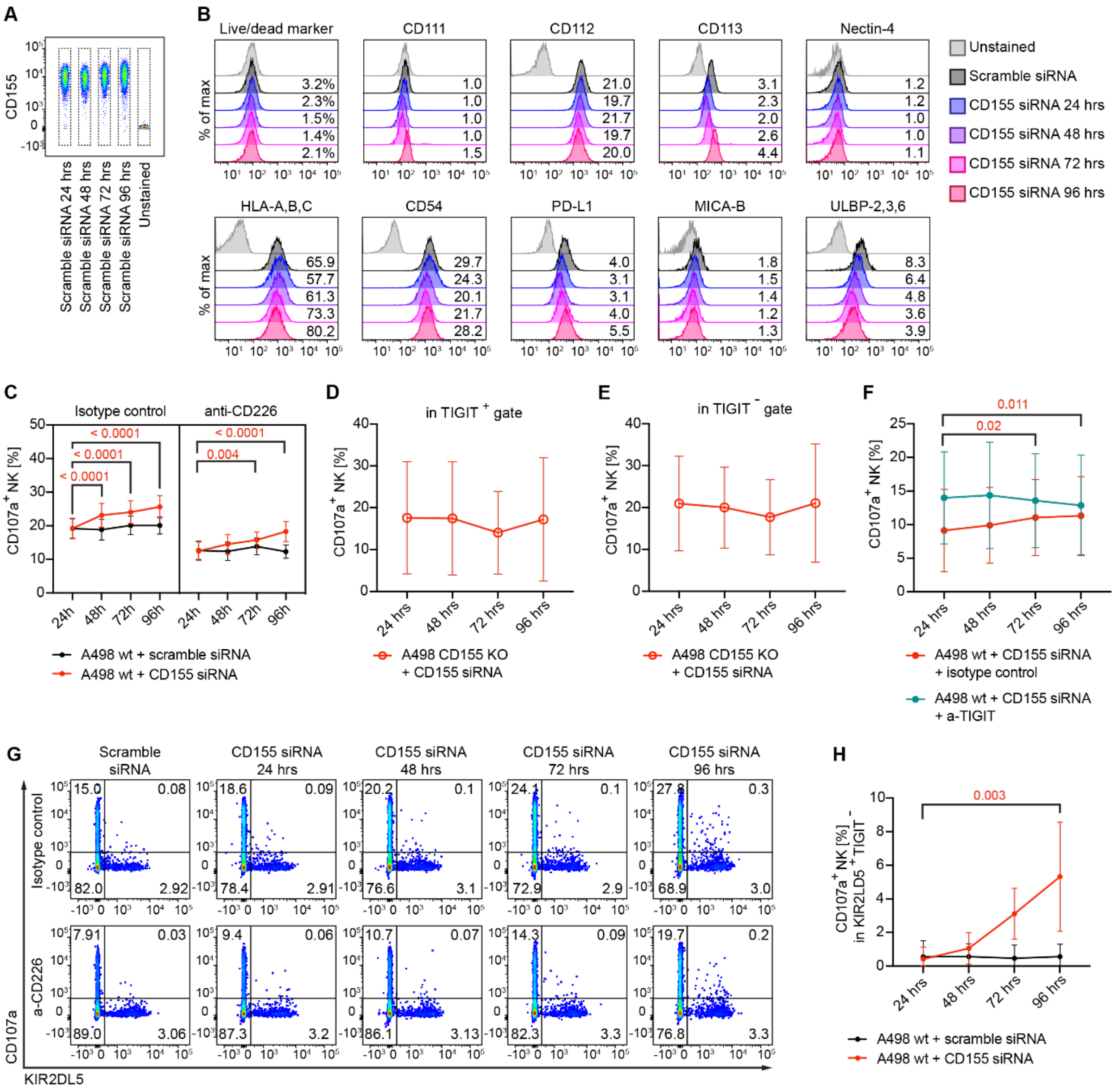


**Fig. S2. Characterization of NK and tumor responses to CD155 silencing on A498 $^{CD155wt\ CD112wt}$ renal carcinoma cells**

**A.** Dot-plot showing the CD155 expression on A498 $^{CD15wt\ CD112wt}$ cells incubated with scramble siRNA for 24, 48, 72 or 96 hours. Data shown are from one representative flow cytometry experiment. **B.** Flow cytometry histograms showing viability and expression of NK ligands on A498 $^{CD15wt\ CD112wt}$ cells incubated with either scramble siRNA (black histogram) or si-RNA CD155 for 24 hours (blue histogram), 48 hours (purple histogram), 72 hours (magenta histogram) or 96 hours (dark red histogram). Light grey histogram: unstained control. Percentage of positive cells (in Live/Dead marker plot) or relative MFI (NK cell ligand plots) are indicated at the right side of the corresponding histogram. No major differences in tumor cell viability and NK ligand expression were detected between conditions. **C.** Three-way plots showing the percentage of CD107a$^+$ NK cells after 2 hours incubation with A498 $^{CD15wt\ CD112wt}$ treated with scramble siRNA (black dots) or CD155 si-RNA (red dots) for 24, 48, 72 or 96 hours, in the presence of isotype control antibody (left panel) or anti-CD226 blocking antibody (right panel). Statistical method: one-way ANOVA followed by Dunnett's Post Hoc test (n=9). The dots localize at the mean values and the lines represent SEM. Mean values with SD are shown in Supporting Table 2. **D, E.** Percentage of degranulating (CD107a$^+$) NK cells for TIGIT$^+$ (**D**) or TIGIT$^-$ (**E**)

NK cells after 2 hours incubation with A498 $^{CD15-/- CD112wt}$ cells treated with CD155 si-RNA for 24, 48, 72 or 96 hours. No statistic has been performed (n=2). Graph shows mean with SEM. **F.** Percentage of degranulating ($CD107a^+$) NK cells after 2 hours incubation with A498 $^{CD15wt CD112wt}$ treated with CD155 si-RNA for 24, 48, 72 or 96 hours, in the presence of an isotype control antibody (red dots) or anti-TIGIT blocking antibody (green dots). Statistical method: two-way ANOVA followed by Dunnett' Post Hoc test (n=3). Graph shows mean with SD. **G.** Representative dot-plots showing NK cell degranulation (CD107a) in relation to KIR2DL5 expression after 2 hours incubation with A498 $^{CD15wt CD112wt}$ treated with scramble siRNA or CD155 si-RNA for 24, 48, 72 or 96 hours (from left to right), in the presence of an isotype control antibody (top row) or anti-CD226 blocking antibody (bottom row). **H.** Percentage of degranulating ($CD107a^+$) NK cells gating on $KIR2DL5^+$ $TIGIT^-$ after 2 hours incubation with A498 $^{CD15wt CD112wt}$ treated with scramble siRNA (black dots) or CD155 si-RNA (red dots) for 24, 48, 72 or 96 hours. Statistical method: two-way ANOVA followed by Dunnett's Post Hoc test (n=3).

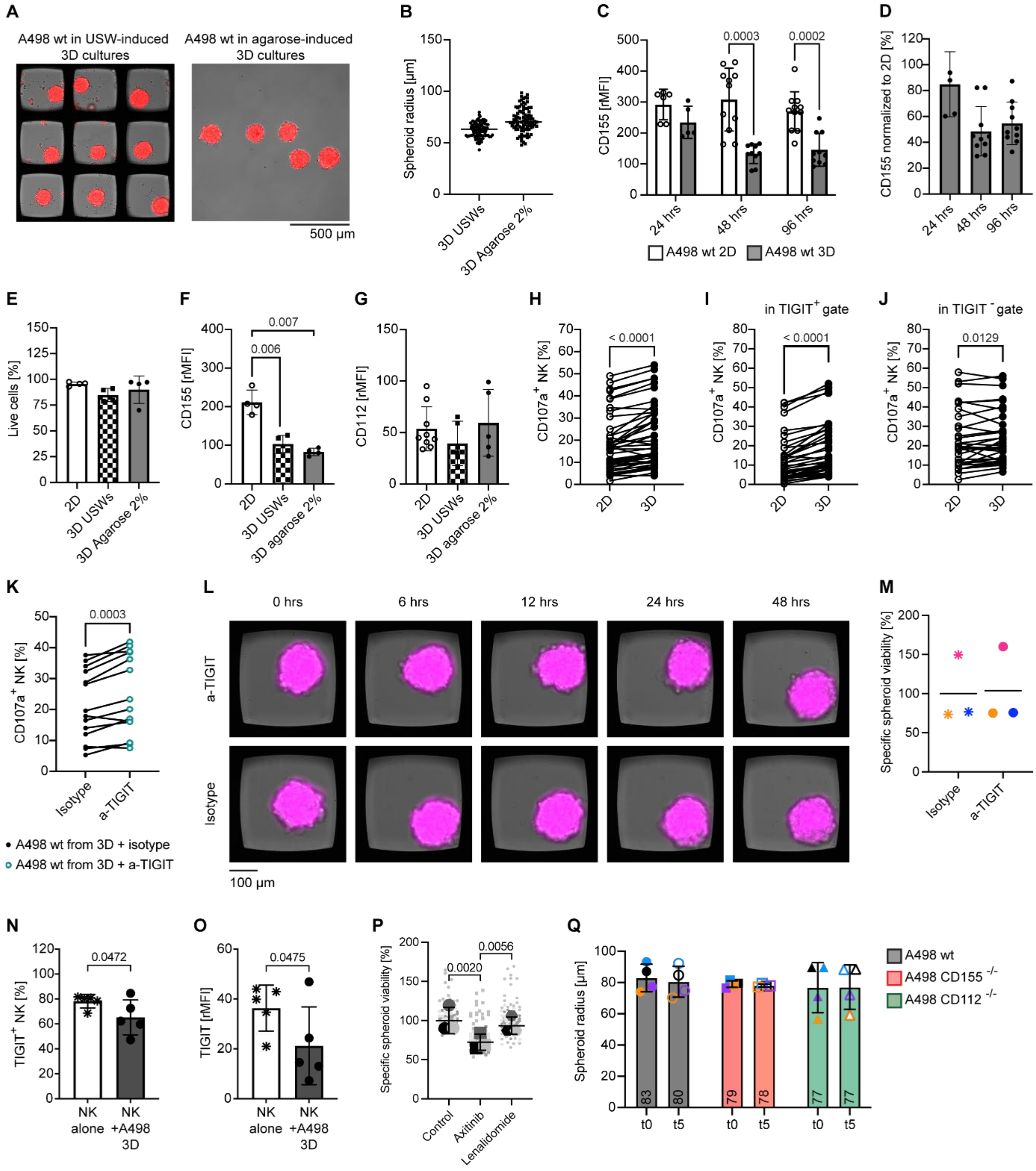

**Fig. S3. Characterization of CD155 and CD112 impact on NK cytotoxic activity and infiltration in solid tumor models with combinatorial treatments**

**A.** Representative wide-field microscopy images of A498-RFP spheroids produced in multichambered microwell chips (left panel) or microwells printed in agarose (right panel). Spheroids were maintained in culture for 48 hours before proceeding with imaging. The thickness of the agarose hydrogels is greater than the working distance of a standard 10x objective, therefore agarose-induced spheroids were transferred into a glass bottom chamber before imaging, while USW-induced spheroids could be imaged directly in the multichambered microwell chip. **B.** Spheroid radius measured from spheroids produced in the multichambered microwell chip (black dots) or agarose microwells (black squares) after 48 hours in culture. Data is from one representative experiment (n=81). Radii were calculated (assuming circular shapes) from areas measured on projected images in ImageJ. **C.** CD155 expression on A498 $^{CD155wt\ CD112wt}$ cells maintained either as 2D cultures (white) or agarose-induced tumor spheroids (dark grey) for 24, 48 or 96 hours. Statistical method: two-way ANOVA (mixed-effect analysis) followed by Tukey's Post Hoc test (24 hours: n=6; 48 hours and 96 hours: n=11). **D.** Percentage of CD155 reduction (normalized for CD155 MFI in 2D cultures at the corresponding timepoint) at 24, 48 or 96 hours. **E-G.** A498 $^{CD155wt\ CD112wt}$ cells were cultured with either standard 2D methods (white-filled bars), agarose-induced 3D culture (black and white-filled bars) or in USW-induced 3D cultures (grey-filled bars) for 48 hours. Flow cytometry was performed on single cell suspensions obtained after enzymatic dissociation to analyze cell viability (**E**) and expression of selected NK cell ligands (**F, G**). Statistical method: one way ANOVA followed by Tukey's Post Hoc test (E, F: n=4; G: n=9). **H.** Pairwise comparison of the percentage of degranulating (CD107a$^+$) NK cells after 2 hours incubation with A498 $^{CD155wt\ CD112wt}$ cells isolated from either 2D cultures or agarose induces-3D cultures. Statistical method: paired t-Test (n=44). **I, J.** Pairwise comparison of the percentage of degranulating (CD107a$^+$) cells among TIGIT$^+$ (**I**) or TIGIT$^-$ (**J**) NK cells after 2 hours incubation with A498 $^{CD155wt\ CD112wt}$ cells isolated from either standard 2D cultures or agarose-induced 3D cultures. Statistical method: paired t-Test (n=37). **K.** Pairwise comparison of degranulating NK cells (CD45$^+$CD107a$^+$ cells) in response to A498 $^{CD155wt\ CD112wt}$ cells isolated from tumor spheroids. In black: isotype control antibody; in teal: anti-TIGIT antibody. Statistical method: paired t-test (n=14). Overnight IL-15 activated NK cells were used as effector cells in H-K. **L.** Time-lapses of A498 $^{CD155wt\ CD112wt}$ spheroid in presence of anti-TIGIT antibody (upper panel) or isotype control antibody (bottom panel). In magenta: TMRM viability dye. **M.** Statistical analysis of specific viability of A498 $^{CD155wt\ CD112wt}$ spheroids in the presence of isotype control (stars) or anti-TIGIT (empty dots) at different timepoints. The big symbols represent the mean values for 36 spheroids; each color represents different replicate. Statistical test: paired t-test (n=3). **N, O.** Expression of TIGIT, shown as percentage of positive cells (**N**) and relative MFI (**O**), analyzed by flow cytometry on NK cells cultured alone (white bar), or as A498 $^{CD155wt\ CD112wt}$ tumor spheroid co-cultures (grey bar). Cells were cultured in the presence of IL-15 for 48 hours at a E:T seeding ratio of 1:2. Statistical method: one way ANOVA followed by Tukey's Post Hoc test (n=7). **P.** Dot-plot showing the specific spheroid viability of individual A498 $^{CD155wt\ CD112wt}$ spheroids (small symbols, n=36) after 12 hours treatment with isotype control antibody (circles), axitinib (squares) or lenalidomide (hexagons). The big symbols represent mean values for 36 spheroids from each experiment. The black bars show the mean value and SD of three experiments (n=3). Statistical method: one-way ANOVA followed by Tukey's Post Hoc test (n=3). **Q.** Radius of A498 $^{CD155wt\ CD112wt}$ (in grey), A498 $^{CD155-/-\ CD112wt}$ (in red) and A498 $^{CD155wt\ CD112-/-}$ (in green) spheroids at timepoint 0 (filled symbols) or after co-culture with IL-15 activated NK cells for 5 hours (empty symbols).

#### Supplementary tables

**Table S1. Percentages of degranulating NK cells gated on TIGIT+ and TIGIT- co-cultured with A498wt cells during CD155-silencing and blocking of CD226 (mean ± standard deviation)**

| | TIGIT+ | | | | TIGIT- | | | |
|---|---|---|---|---|---|---|---|---|
| | Scramble Si-RNA | | CD155 Si-RNA | | Scramble Si-RNA | | CD155 Si-RNA | |
| Time (h) | IgG | a-CD226 | IgG | a-CD226 | IgG | a-CD226 | IgG | a-CD226 |
| 24 | 12.8 ± 8.4 | 7.9 ±6.5 | 14.2 ± 9.8 | 8.6 ± 7.1 | 25.0 ± 9.6 | 18.1 ± 9.3 | 24.0 ± 10.6 | 18.1 ± 9.4; |
| 48 | 12.8 ± 8.9 | 7.6 ± 6.6 | 19.9 ± 11.5 | 11.7 ± 8.7 | 24.7 ± 9.8 | 18.1 ± 8.9 | 27.0 ± 11.1 | 19.2 ± 8.8; |
| 72 | 14.6 ± 8.4 | 9.1 ± 6.4 | 21.5 ± 10.8 | 13.7 ± 7.9; | 25.4 ± 8.8 | 19.4 ± 8.3 | 26.4 ± 9.7 | 18.7 ± 7.7; |
| 96 | 14.7 ± 7.8 | 8.0 ± 5.6 | 24.0 ± 10.8 | 16.5 ± 10.4 | 25.2 ± 8.9 | 17.7 ± 6.5 | 26.7 ± 8.1. | 20.7 ± 8.1. |

**Table S2. Percentages of total degranulating NK cells co-cultured with A498wt cells during CD155-silencing and blocking of CD226 (mean ± standard deviation)**

| | Scramble Si-RNA | | CD155 Si-RNA | |
|---|---|---|---|---|
| Time (h) | IgG | a-CD226 | IgG | a-CD226 |
| 24 | 19.2 ± 8.7 | 12.7 ± 7.8 | 19.2 ± 11.0 | 12.5 ± 7.7 |
| 48 | 18.9 ± 9.2 | 12.4 ± 7.8 | 23.1 ± 12.8 | 14.5 ± 8.3 |
| 72 | 20.1 ± 8.2 | 13.9 ± 7.1 | 24.1 ± 12.1 | 15.8 ± 7.0 |
| 96 | 20.1 ± 7.6 | 12.3 ± 5.6 | 25.6 ± 12.1 | 18.3 ± 8.4 |
